# Brief temporal perturbations in somatosensory reafference disrupt perceptual and neural attenuation and increase supplementary motor-cerebellar connectivity

**DOI:** 10.1101/2022.11.25.517892

**Authors:** Konstantina Kilteni, Christian Houborg, H. Henrik Ehrsson

## Abstract

Intrinsic delays in sensory feedback can be detrimental for motor control. As a compensation strategy, the brain predicts the sensory consequences of movement via a forward model on the basis of a copy of the motor command. Using these predictions, the brain attenuates the somatosensory reafference to facilitate the processing of exafferent information. Theoretically, this predictive attenuation gets disrupted by (even minimal) temporal errors between the predicted and the actual reafference, but direct evidence for such disruption is lacking since previous neuroimaging studies contrasted conditions of nondelayed reafferent input with exafferent one. Here, we combined psychophysics with functional magnetic resonance imaging to test whether subtle perturbations in the timing of somatosensory reafference disrupt its predictive processing. Twenty-eight participants generated touches on their left index finger by tapping a sensor with their right index finger. The touches on the left index finger were delivered at the time of the two fingers’ contact or with a 100 ms delay. We found that such brief temporal perturbations disrupted the attenuation of the somatosensory reafference both at the perceptual and neural level, leading to greater somatosensory and cerebellar responses and weaker somatosensory connectivity with the cerebellum proportionally to perceptual changes. Moreover, we observed increased connectivity of the supplementary motor area with the cerebellum during the perturbations. We interpret these effects as the failure of the forward model to predictively attenuate the delayed somatosensory reafference and the return of the prediction error to the motor centers, respectively.

**Significance statement:** Our brain receives the somatosensory feedback of our movements with delay. To counteract these delays, motor control theories postulate that the brain predicts the timing of the somatosensory consequences of our movements and attenuates sensations received at that timing. This makes a self-generated touch feel weaker than an identical external touch. However, how subtle temporal errors between the predicted and the actual somatosensory feedback perturb this predictive attenuation remains unknown. We show that such errors make the otherwise attenuated touch feel stronger, elicit stronger somatosensory responses, weaken the cerebellar connectivity with somatosensory areas, and increase it with motor areas. These findings show that motor and cerebellar areas are fundamental in forming temporal predictions about the sensory consequences of our movements.

## Introduction

During voluntary movement, our sensorimotor loop suffers from ubiquitous delays due to sensory transduction, neural conduction and brain processing of the sensory feedback (Wolpert and Flanagan, 2001; Franklin and Wolpert, 2011). These delays have a non-negligible magnitude even exceeding ∼100 ms (Scott, 2016), and their impact can be detrimental through destabilizing our motor output and leading to oscillatory movements when rapidly correcting motor errors (Miall and Wolpert, 1996; Kawato, 1999). To compensate for the delayed feedback, the brain uses a forward model in combination with a copy of the motor command (*efference copy*) to predict the sensory consequences of the movement and thus, rely less on the delayed input (Shadmehr et al., 2010; McNamee and Wolpert, 2019). These predictions allow to prospectively correct the motor command in case of errors (Shadmehr et al., 2010), and improve the estimation of the current state of our body (Todorov and Jordan, 2002; Scott, 2004; Shadmehr et al., 2008).

The forward model-based predictions further serve to differentiate sensory reafference from exafference. Both animal and human research has repeatedly shown that signals received at the predicted time – and thus corresponding to the sensory consequences of the movement – are suppressed, to facilitate the processing of external signals (Blakemore et al., 2000b; Brooks and Cullen, 2019; McNamee and Wolpert, 2019; Audette et al., 2021). For example, when moving the right hand to touch the left hand, the reafferent touches on the left hand feel systematically weaker (Blakemore et al., 1999; Shergill et al., 2003; Kilteni and Ehrsson, 2017a, 2017b, 2022; Kilteni et al., 2018, 2020; Asimakidou et al., 2022) and elicit weaker somatosensory responses compared to exafferent touches of identical intensity (Blakemore et al., 1998; Hesse et al., 2010; Kilteni and Ehrsson, 2020). Critically, this attenuation of sensory reafference is time-locked to the expected feedback time and it is reduced, or even vanished, when identical somatosensory input is presented at close temporal proximity, either earlier (Bays et al., 2005) or later (Blakemore et al., 1999; Bays et al., 2005; Kilteni et al., 2019, 2021).

From a theoretical perspective, the cerebellum implements the forward model and predicts the sensory consequences of the movement (Shadmehr et al., 2008; McNamee and Wolpert, 2019; Popa and Ebner, 2019) using the efference copy provided by the supplementary motor area (Haggard and Whitford, 2004; Pynn and DeSouza, 2013) to attenuate the reafferent somatosensory input. These computational processes are very sensitive to errors between the predicted and the actual sensory feedback (Wolpert and Flanagan, 2001; Shadmehr et al., 2010): under certain conditions, errors can force the sensorimotor system to either refine its motor plan (Johnson et al., 2019), re-optimize the forward model’s predictions after systematic exposure to the errors (Izawa et al., 2008), or disregard them and attribute them to external causes when they are large (Wei and Körding, 2009; Wilke et al., 2013). However, previous human neuroimaging studies manipulated the timing of somatosensory feedback imposing large perturbations (reaching 400-500 ms) (Blakemore et al., 2001; Shergill et al., 2013), effectively contrasting conditions of nondelayed reafferent with rather exafferent input. Thus, how subtle temporal perturbations in somatosensory reafference disrupt its predictive processing remains unknown.

By combining psychophysics with functional magnetic resonance imaging (fMRI), we investigated perceptual and neural responses to the presence (50% trials) or absence (50% trials) of brief temporal perturbations (100 ms) between the participants’ right hand movements and the somatosensory feedback on their left hand. Such brief delays are not typically detectable (Blakemore et al., 1999) and do not lead to sensorimotor adaptation if non-persistent (Kilteni et al., 2019). However, they should theoretically disrupt the sensorimotor loop in two ways. First, delays should interrupt the attenuation of the somatosensory reafference by the forward model, leading to greater somatosensory and cerebellar responses, and weaker somatosensory connectivity with the cerebellum. Second, they should increase the connectivity of the supplementary motor area with the cerebellum expressing the conveyance of the error to the motor centers.

## Materials and Methods

### Participants

After providing written informed consent, twenty-nine (29) volunteers (15 women, 14 men; 27 right-handed, 2 ambidextrous) aged 19-38 years old participated in the study. Handedness was assessed using the Edinburgh Handedness Inventory (Oldfield, 1971). The sample size was set to 30 based on our previous study (Kilteni and Ehrsson, 2020), but due to scanner technical issues, fMRI data were collected from 29 individuals. After data collection, one participant was further excluded, for giving the same response to almost all trials (49 out of 50 trials) of one of the two conditions of the psychophysical task, making the psychophysical modeling unreliable. To be consistent, this participant was excluded also in the fMRI analysis. Therefore, both behavioral and fMRI analyses were performed with a total of 28 participants (14 women, 14 men; 26 right-handed, 2 ambidextrous; 19-38 years old).

### Psychophysics and fMRI

The fMRI scan was conducted before the psychophysics session for practical reasons. The psychophysics experiment was conducted in the MR scanner environment using the same equipment (same motor setup and force sensors) as used in the fMRI session (see further below). After the fMRI experiment and the psychophysics session, additional fMRI runs and psychophysical tasks were conducted as part of a different study addressing a separate question, which we do not report in the current manuscript. The Ethics Review Authority approved the study (project: #2016/445-31/2, amendment: #2018:1397-32).

### Procedures and experimental design for the psychophysical task

The psychophysical task was a two-alternative forced-choice force-discrimination task (**Figure 1a**) that has been extensively used to assess somatosensory attenuation in previous studies (Bays et al., 2005, 2006; Kilteni et al., 2019, 2020, 2021; Asimakidou et al., 2022; Kilteni and Ehrsson, 2022), and it served to quantify the perceived intensity of *nondelayed* (0 ms) and *delayed* (100 ms) self-generated touches. Participants laid comfortably in a supine position on the MRI scanner bed. Their left hands were placed palm-up on an MR-compatible plastic table with their left index finger in contact with a 3D-printed probe that contained a force sensor and that was controlled by a motor through string-based transmission. Their right index finger was placed next to a second force sensor that was also placed on the table, on top of (but not in contact with) the probe on the left index finger (**Figure 1b**). Both arms were supported by sponges to maximize the comfort of the participants.

**Figure 1.**
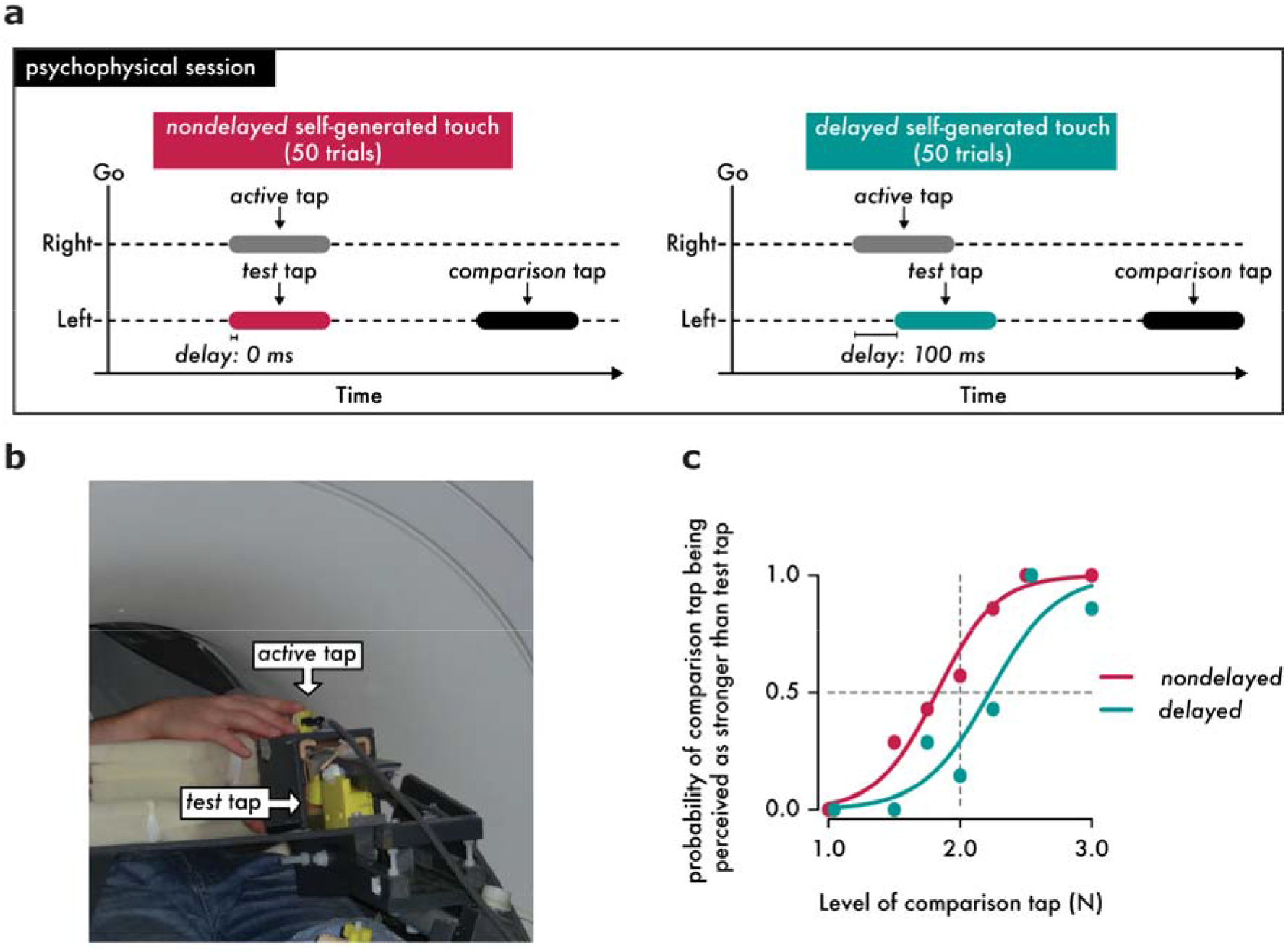
The psychophysical session conducted inside the MR scanner. **(a)** Participants performed a force discrimination task to assess the participants’ perceived magnitude of nondelayed and delayed self-generated touches of 250 ms duration. In this task they received two taps (the *test* and the *comparison* tap) on the pulp of their left index fingers from an electric motor. The participants produced the *test* tap on their left index finger by actively tapping a sensor with their right index finger (gray rectangle, *active* tap) and received the *test* tap with a 0 ms (magenta rectangle, left) or a 100 ms delay (cyan rectangle, right). Following, a second tap (*comparison* tap) was applied to their finger with a variable magnitude (black rectangle), and participants verbally reported which of the two taps applied on their left index finger (*i*.*e*., the *test* or the *comparison* tap) felt stronger. **(b)** Overview of the fMRI-compatible setup used in the psychophysics experiments and in the fMRI experiment. The psychophysics task was performed while the subjects were laying on the scanner bed but without being scanned. **(c)** Responses and fitted logistic models of the responses of one example participant in the two experimental conditions. Two data points are horizontally jittered to avoid complete overlap.

During the task, the participants were asked to tap with their right index finger the force sensor (*active tap*) after an auditory Go cue. Prior to the task we instructed the participants to tap the sensor with their right index finger at an intensity that is comfortable for them and to keep the same style of taps throughout the session. The *active* tap of the right index finger (force exceeded > 0.4 N) was used to trigger the *test tap* on their left index finger after a 0 ms or a 100 ms. The *test* tap had a fixed intensity of 2 N. The intrinsic delay of the system (*i*.*e*., time difference between the *active* tap exceeding 0.4 N until the *test* tap reaches 80% of its maximum magnitude) was ∼ 53 ms. After a random delay between 800 and 1500 ms, participants received a subsequent externally generated tap (*comparison* tap) of variable intensity (1, 1.5, 1.75, 2, 2.25, 2.5, or 3 N). The taps were applied for approximately 250 ms (mean ± s.e.m.: 249.209 ± 5.146 ms). Participants were asked to verbally indicate which tap (the *test* or the *comparison* tap) felt stronger on their left index finger. Each level of the *comparison* tap was repeated 7 times, except for the level of 2 N that was repeated 8 times. Consequently, each condition consisted of 50 trials, resulting in 100 trials per participant. The order of conditions was randomized across participants. On average, participants pressed an *active* tap of (mean ± s.e.m.) 2.328 ± 0.203 N with their right index finger and received a *test* tap of 1.997 ± 0.004 N on their left index finger. The mean duration of the *active* tap produced by the participants was ∼180 ms (mean ± s.e.m.: 176.432 ± 10.888 ms) while the duration of the *test* tap produced by the setup was 250 ms, as mentioned earlier.

### Processing, hypotheses, and statistical analysis of psychophysical data

There were no missing trials from any participant in any of the two conditions, resulting to a total of 2800 trials (28*50*2 = 2800 trials). After data collection, we excluded any psychophysical trials in which the participants did not tap the sensor with their right index finger after the GO cue, tapped lightly and did not trigger the touch on the left index finger (*active* tap < 0.4. N), tapped more than once, tapped before the GO cue, or any trials in which the *test* tap was not applied correctly (*test* tap <1.85 N or *test* tap > 2.15 N). This resulted to the exclusion of 117 trials out of 2800 psychophysical trials (4.18%).

We fitted the participants’ responses with a generalized linear model (**Figure 1c**), using a *logit* link function (Equation 1):

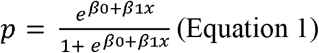

We extracted two parameters of interest: the Point of Subjective Equality 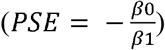, which represents the intensity at which the *test* tap felt as strong as the *comparison* tap (p = 0.5) and quantifies the perceived intensity of the *test* tap, and the Just Noticeable Difference 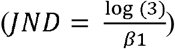 which reflects the participants’ discrimination capacity. The PSE and JND are independent qualities of sensory judgments: higher PSE values indicate a stronger perceived magnitude, while higher JND values indicate a lower force discrimination capacity (*i*.*e*., lower somatosensory precision). To quantify the difference in the perceived magnitude of the self-generated touch in the two conditions, we calculated the difference between the PSEs of the two conditions (*PSE*_*delayed*_ – *PSE*_*nondelayed*_).

Based on previous studies (Blakemore et al., 1999; Bays et al., 2005; Kilteni et al., 2019, 2021), we hypothesized a significant difference between the PSEs of the two conditions, with the *delayed self-generated touch* condition yielding a greater magnitude of the perceived touch due to the temporal perturbation compared to the *nondelayed self-generated touch* condition. We hypothesized no differences in the discrimination capacity (JND) between the two conditions, given our previous results involving the same right index finger movement and touch on the left index finger (Asimakidou et al., 2022; Kilteni and Ehrsson, 2022). Psychophysical data were analyzed using R (2022) and JASP (2022). Data normality was assessed using the Shapiro–Wilk test, and planned comparisons were made using parametric (paired *t*-test) statistical tests given that the data were normally distributed. For each test, 95% confidence intervals (*CI*^*95*^) are reported. Effect sizes are given by Cohen’s *d*. A Bayesian factor analysis was carried out for non-significant statistical comparisons of interest (default Cauchy priors with a scale of 0.707) to provide information about the level of support for the null hypothesis compared to the alternative hypothesis (*BF*_*01*_). Correlations between perceptual and neural responses (see below) were assessed with the Kendall (*tau-b*) or Pearson (ρ) correlation coefficients depending on the data normality. All statistical tests were two-tailed.

### Complementary post-hoc psychophysical analysis

We performed a control analysis to test for the absence of any significant learning effects due to repeated exposure to the 100 ms delay in the *delayed self-generated touch* condition. According to one of our previous study (Kilteni et al., 2019), significant learning of a 100 ms delay requires more than 400 exposure trials (50 initial exposure trials and 350 re-exposure trials). Here, participants were exposed to only 50 trials during the psychophysical assessment and thus no learning should be observed. However, if there is indeed some adaptation, this may reduce the effect of the brief temporal perturbations on the psychophysical responses especially at the end of the psychophysical task. To confirm the absence of such adaptation to delays, we fitted the participants responses in the *nondelayed* and *delayed self-generated touch* condition separately for the first and second half of the task and we compared the difference in PSEs between the two halves using a paired *t*-test given that the data were normally distributed.

### Procedures and experimental design for the fMRI experiment

The fMRI session always preceded the force-discrimination task for practical reasons. Using the same equipment and identically to the psychophysical session, the participants were asked to tap the force sensor with their right index finger (*active* tap) after the auditory GO cue and received the *test* tap on their left index finger (2 N), with or without the 100 ms delay. Blocks including 24 such trials (with or without the delay) were interleaved with rest blocks of 16 seconds during which the subjects remained relaxed (**Figure 2**). We chose alternating blocks of only 24 trials to avoid learning of a 100 ms delay due to repeated exposure to the delay, given our previous study (Kilteni et al., 2019) showing that more than 400 exposure trials are needed for participants to adapt to a 100 ms sensorimotor delay. Messages were displayed on a screen seen through a mirror attached to the head coil and informed the participants about what they had to do (‘PRESS or ‘PAUSE). Participants were asked to fixate their gaze on the fixation cross seen on the screen and follow the messages. The participants’ right arm and hand were peripherally visible. There were 12 blocks of self-generated touches (6 with and 6 without delay) and 12 blocks of rest, resulting to 144 nondelayed and 144 delayed self-generated touch trials. The condition blocks were alternating, and their order was randomized between participants. On average, participants pressed an *active* tap of (mean ± s.e.m.) 2.084 ± 0.236 N with their right index finger and received a *test* tap of 1.996 ± 0.007 N on their left index finger. As in the psychophysical task, the mean duration of the *active* tap produced by the participants was ∼180 ms (mean ± s.e.m.: 176.049 ± 9.416 ms) while the duration of the *test* tap produced by the setup was ∼250 ms (mean ± s.e.m.: 241.175 ± 4.604 ms).

**Figure 2.**
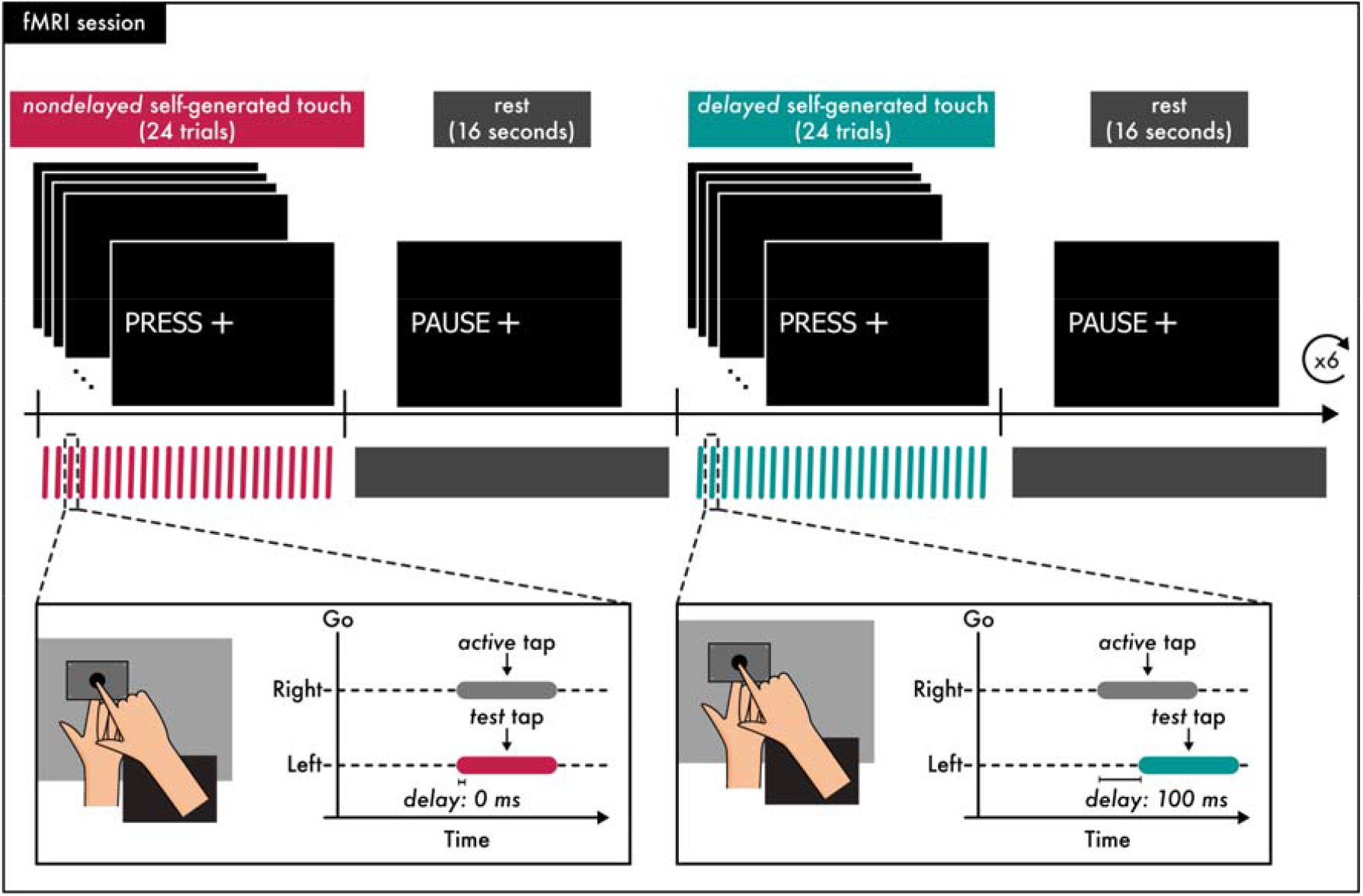
The fMRI session. The functional run was organized in blocks during which the participants produced a self-generated touch and blocks in which they remained relaxed. The run began with a block in which participants received the message “PRESS” on the screen that instructed them to tap the force sensor with their right hand (*active* tap) and received the *test* tap on their left index finger (2 N, with or without the 100 ms delay). Participants were required to perform 24 such trials. In the next block participants received the message “PAUSE” which instructed them to relax both their hands for 16 seconds. The next block required again participants to produce 24 self-generated taps (with or without a delay), followed by a rest block of 16 seconds. Each block of self-generated touches (with or without a delay) was repeated 6 times and blocks of the different conditions were alternating. The proportions of *nondelayed* and *delayed* self-generated touches were equal (50%).

### Preprocessing, hypotheses, and primary statistical analysis of fMRI activations

fMRI acquisition was performed using a General Electric 3T scanner (GE750 3T) equipped with an 8-channel head coil. T2-weighted echo-planar images (EPIs) containing 42 slices were acquired (repetition time: 2000 ms; echo time: 30 ms; flip angle: 80°; slice thickness: 3 mm; slice spacing: 3.5 mm; matrix size: 76 × 76; in-plane voxel resolution: 3 mm). A total of 330 functional volumes were collected for each participant. For the anatomical localization of activations, a high-resolution structural image containing 180 slices was acquired for each participant before the acquisition of the functional volumes (repetition time: 6404 ms; echo time: 2.808 ms; flip angle: 12°; slice thickness: 1 mm; slice spacing: 1 mm; matrix size: 256 × 256; voxel size: 1 mm x 1 mm x 1 mm).

We ran a standard preprocessing pipeline using the CONN toolbox (version 21a) (Whitfield-Gabrieli and Nieto-Castanon, 2012) including realignment, unwarping and slice-time correction. Outlier volumes were detected using the Artifact Detection Tools employing the option for liberal thresholds (global-signal threshold of *z* = 9 and subject-motion threshold of 2 mm). Following, we simultaneously segmented the images into gray matter, white matter and cerebrospinal fluid and normalized into standard MNI space (Montreal Neurological Institute, Canada). Next, the images were spatially smoothed using an 8 mm FWHM Gaussian kernel. The structural images were also simultaneously segmented (into gray and white matter and cerebrospinal fluid) and normalized to MNI space.

The preprocessed data were analyzed with a general linear model (GLM) for each participant in Statistical Parametric Mapping 12 (SPM12; Welcome Department of Cognitive Neurology, London, UK, http://www.fil.ion.ucl.ac.uk/spm). We used an event-related design with trial onsets defined as the timings when the magnitude of the *test* tap peaked, and zero trial durations. Regressors of interest were included for each of the two conditions of interest (*delayed* and *nondelayed self-generated touch*). Similar to the psychophysical session, any trials in which the participants did not tap the sensor with their right index finger after the auditory cue, tapped but did not trigger the touch on the left index finger (*active* tap < 0.4N), tapped more than once, or tapped before the auditory GO cue, were excluded from the regressors of interest and modelled as four (4) individual regressors of no interest. This resulted to the exclusion of 119 trials out of 8064 fMRI trials from the main regressors (1.48%). In addition, the six motion parameters, and any outlier volumes were included as regressors of no interest. The trials of each condition were convolved with the canonical hemodynamic response function of SPM 12. The first level analysis was restricted to gray matter voxels using a binary (threshold 0.2) and smoothed mask (8 mm FWHM Gaussian kernel) of gray matter, that was based on the individual’s segmented structural image (gray matter). Contrasts between the two condition regressors of interest (*delayed* > *nondelayed* and *nondelayed* > *delayed*) were created. At the second level of analysis, random-effects group analyses were performed by entering the contrast images from each subject into a one sample t-test. Contrasts of interest focused on the comparison *delayed* > *nondelayed* and *nondelayed* > *delayed*.

We hypothesized that the activity of the right somatosensory cortices will differ between the *nondelayed* and *delayed* self-generated touch conditions. To correct for multiple comparisons in right somatosensory areas, we performed small volume corrections within spherical regions of interest (ROIs) of 10-mm radius, centered at peaks detected in our previous study using the same scanner, same equipment and same tactile stimulation (2 N) on the same finger (left index finger) (Kilteni and Ehrsson, 2020). These peaks corresponded to the right primary somatosensory cortex (right S1) (MNI: *x* = 50, *y* = −20, *z* = 60), and the right secondary somatosensory cortex (rSII)) (MNI: *x* = 46, *y* = −14, *z* = 16). To correct for multiple comparisons within the cerebellum, we used anatomical masks created with the Anatomy toolbox (Eickhoff et al., 2005) including the hemispheres of the right and left lobules V, VI and VIII, given the involvement of these cerebellar regions in the sensorimotor cerebellar body representation (Grodd et al., 2001; Diedrichsen et al., 2005; Stoodley and Schmahmann, 2009; O’Reilly et al., 2010; Buckner et al., 2011; Bostan et al., 2013; Guell et al., 2018; King et al., 2018). To directly compare our results to those from the study of Blakemore (2001) using Positron Emission Tomography (PET), we also included a mask containing the right lobule VIIa Crus I given that the authors reported peaks in both lobules VI and VIIa Crus I.

For each peak activation, the coordinates in MNI space, the *z* value and the *p* value are reported. We denote that a peak survived a threshold of *p* < 0.05 after correction for multiple comparisons at the whole-brain or small volume by the term “*FWE-corrected*” following the *p* value.

### Statistical analysis of the relationship between fMRI activations (*delayed* > *nondelayed* self-generated touch) and the psychophysical results

We tested for a relationship between the perceptual differences in force discrimination revealed by the psychophysical task and the effects revealed by our fMRI univariate analysis. To do so, we extracted the signal from the contrast estimates of each condition against zero (*nondelayed* > 0 and *delayed* > 0) using the Marsbar Toolbox (Brett et al., 2002), at the peaks where the activity significantly differed between the two conditions *(p < 0*.*05 FWE-corrected)*. We then performed a standard correlation analysis between the signal difference between the two conditions at the significant peaks and the difference in the PSEs extracted from the psychophysical task (*PSE*_*delayed*_ – *PSE*_*nondelayed*_).

### fMRI functional connectivity: preprocessing, hypotheses, and statistical analysis

For the functional connectivity analysis, data were further denoised using the component-based noise correction method (*CompCor*) as it is implemented in the CONN toolbox. Five principal components from white matter, five principal components from cerebrospinal fluid, twelve principal realignment components (six plus 1^st^ order derivatives) and scrubbing parameters, together with two principal components per condition (the time series and its first derivative), were extracted and used as confounds. A bandpass filter [0.008 Hz, Inf] was applied, and the data were linearly detrended.

We previously showed that the degree of functional connectivity between the cerebellum and the somatosensory areas is linearly and positively related to the degree to which participants perceptually attenuated their self-generated touches (Kilteni and Ehrsson, 2020). Therefore, we hypothesized that the right somatosensory cortices would *decrease* their connectivity with the cerebellum when a temporal perturbation is present as a function of the participants’ perception. To test this hypothesis, we conducted a seed-to-voxel analysis in the form of generalized psychophysiological interactions (*gPPI*) (McLaren et al., 2012) using the denoised data. Right somatosensory seeds of interest were defined as spheres with a 8-mm radius around the two somatosensory peaks (right S1 and right SII, *p* < 0.05 *FWE-corrected*) revealed by the activation analysis (*delayed* > *nondelayed* self-generated touches) of the present study. At the group level, the contrasts of interest consisted of the effect of delay (*delayed* > *nondelayed*) and we identified both increases and decreases in the functional connectivity of the seeds. To specifically identify any connectivity changes of the somatosensory seeds that scaled with the participants’ perception, we used the PSE difference from the psychophysical task as a second-level covariate (*PSE*_*delayed*_ – *PSE*_*nondelayed*_).

Given that the supplementary motor area is theorized to provide the efference copy to predict and attenuate self-generated somatosensory activity (Haggard and Whitford, 2004) but also to use information related to discrepancies between the predicted and the actual feedback to update the motor plan (Welniarz et al., 2021), we further hypothesized connectivity changes between conditions with the left supplementary motor area (left SMA). Specifically, we anticipated that the left SMA will increase its connectivity with the left cerebellum (left CB) in presence of the temporal perturbations due to the feedback signal indicating the temporal discrepancy between the predicted and the actual touch on the left index finger. At the same time, the left SMA should decrease its connectivity with the right somatosensory cortices during the temporal perturbations, indicating the reduced attenuation of the somatosensory reafference on the left hand, similar to our hypothesis about the cerebellum. To test the left SMA connectivity, we placed a seed of interest (8-mm radius sphere) at the peak corresponding to the left supplementary motor area that showed significant activation in both condition contrasts against zero (*p* < 0.05 *FWE-corrected*). Since we did not have a hypothesis whether these theorized effects will be mediated by the participants’ somatosensory perception, we performed two connectivity analyses, with and without the participants’ perceptual changes (*PSE*_*delayed*_ – *PSE*_*nondelayed*_) as a covariate.

Statistical maps were assessed using corrections for multiple comparisons using either anatomical masks or peaks from our previous study (Kilteni and Ehrsson, 2020). When using the somatosensory seeds (right S1, right SII), we corrected for multiple comparisons within the cerebellum, by performing small volume corrections within anatomical masks (right and left lobules V, VI and VIII), identically to our univariate analysis. To correct for the left SMA, we used a spherical ROI (10-mm radius) around the left supplementary motor area peak detected in our previous study (MNI: *x* = −6, *y* = −8, *z* = 54) (Kilteni and Ehrsson, 2020). When using the left SMA as seed, we corrected for somatosensory areas by performing small volume corrections within the spherical ROIs (10-mm radius) centered at the two peaks (right S1, right SII) detected in our previous study (Kilteni and Ehrsson, 2020), identically to our univariate analysis. Corrections for multiple comparisons within the cerebellum were performed using the above-mentioned masks.

### Complementary post-hoc fMRI analyses

In a subsequent analysis, we explored the potential influence of small variations in magnitude of the self-generated force in *active* taps on the BOLD signal. We know that larger muscular forces can produce increased BOLD signal in the primary motor cortex, posterior supplementary motor area, cerebellum, and secondary somatosensory cortex (Dettmers et al., 1995; Ehrsson et al., 2001), although these previous studies used much larger force variations compared to the small variations expected in the current study. We followed the same modelling approach described above, but we also included the magnitude of the *active* tap on each trial as a parametric modulator for all the trials of the two conditions of interest. The two contrasts of interest focused on the overall modulation of the *active* taps across both conditions (*pmod*_*delayed*_ + *pmod*_*nondelayed*_ > 0) and the effect of delay (*delayed* > *nondelayed*). We expected that the left motor cortex (left M1) and the right cerebellum (right CB) might increase their activity as a function of the magnitude of the *active* tap – that is, stronger forces of the right hand will elicit stronger motor activity in the left hemisphere and stronger cerebellar activity in the right hemisphere. To test for these hypothesis, we performed small volume corrections within a spherical ROI centered at the left primary motor cortex (MNI: *x* = −38, *y* = −12, *z* = 52) detected in our previous study (Kilteni and Ehrsson, 2020) and within the above-mentioned anatomical cerebellar masks. Then we conducted an additional control analysis for the condition-specific (*delayed* > *nondelayed* and *nondelayed* > *delayed)* by including the force parametric modulator in the model and regressing out force-related signal variation in the data.

Similar to our control analysis in the psychophysics task, we performed a control analysis to test for the absence of any significant learning effects due to repeated exposure to the 100 ms delay. As mentioned above, significant learning of a 100 ms delay requires more than 400 exposure trials (Kilteni et al., 2019). To avoid this and elucidate genuine differences between the *nondelayed* and *delayed* self-generated touch conditions, we thus designed the run to include only 24 trials on each block, with blocks being constantly alternating. However, if there is indeed some adaptation, this may reduce the effect of the brief temporal perturbations on the BOLD signal especially towards the ends of the fMRI run. To confirm the absence of such adaptation to delays, we modeled the trials of each condition (*nondelayed* and *delayed* self-generated touch) separately for each block (1^st^, 2^nd^, 3^rd^, 4^th^, 5^th^, and 6^th^) resulting to 12 different regressors. We then created contrasts of each condition against zero for the first and the last block (*e*.*g*., *nondelayed*_block1_ > 0, *delayed*_block1_ > 0, *nondelayed*_block6_ > 0, *delayed*_block6_ > 0) and we extracted the activity from each contrast at the peak voxels revealed by the univariate analysis (across all blocks) using the Marsbar Toolbox (Brett et al., 2002). We then performed a paired *t*-test with the difference between the two conditions between the first and the last block given that the data were normally distributed.

## Results

### Temporal perturbations disrupt the perceptual attenuation of somatosensory reafference

For all participants and all conditions, the fitted logistic models were very good, with McFadden’s R squared measures ranging between 0.409 and 0.945 (**Supplementary Figure S1)**. The *nondelayed self-generated touch* condition produced a significant decrease in the PSE (*i*.*e*., attenuation) compared to the *delayed self-generated touch* condition (*n* = 28, *t*(27) *=* −5.726, *p* < 0.001, *Cohen’s d* = −1.082, *CI*^*95*^ = [−0.297, −0.140]) despite having identical intensities (*i*.*e*., 2 N) (**Figure 3a**). This effect was observed for 23 out of 28 participants (82.1%). Together, these findings replicate previous results (Blakemore et al., 1999; Bays et al., 2005; Kilteni et al., 2019, 2021) showing that a self-generated touch feels stronger when delivered with a 100 ms delay compared to an identical self-generated touch delivered at the time of contact between the two fingers. Moreover, there was no difference in the force discrimination capacity (JND) between the two conditions (*n* = 28, *t*(27) *=* −1.048, *p* = 0.304, *Cohen’s d* = −0.198, *CI*^*95*^ = [−0.068, 0.022]) (**Figure 3b**). Specifically, 14 participants (50%) increased their JNDs, and 14 participants (50%) decreased their JNDs between conditions. A Bayesian analysis also supported the absence of a JND difference (*BF*_*01*_ = 3.033). Together, the psychophysical results indicate that the attenuation of somatosensory reafference observed when the touch is delivered at its expected time (*nondelayed*) (PSE) gets disrupted when the same touch is delivered with a delay (*i*.*e*., 100 ms) but without influencing the somatosensory precision (JND) (Asimakidou et al., 2022; Kilteni and Ehrsson, 2022). Thus, collectively the psychophysical results corroborated that our behavioral paradigm worked as expected in the scanner environment.

**Figure 3.**
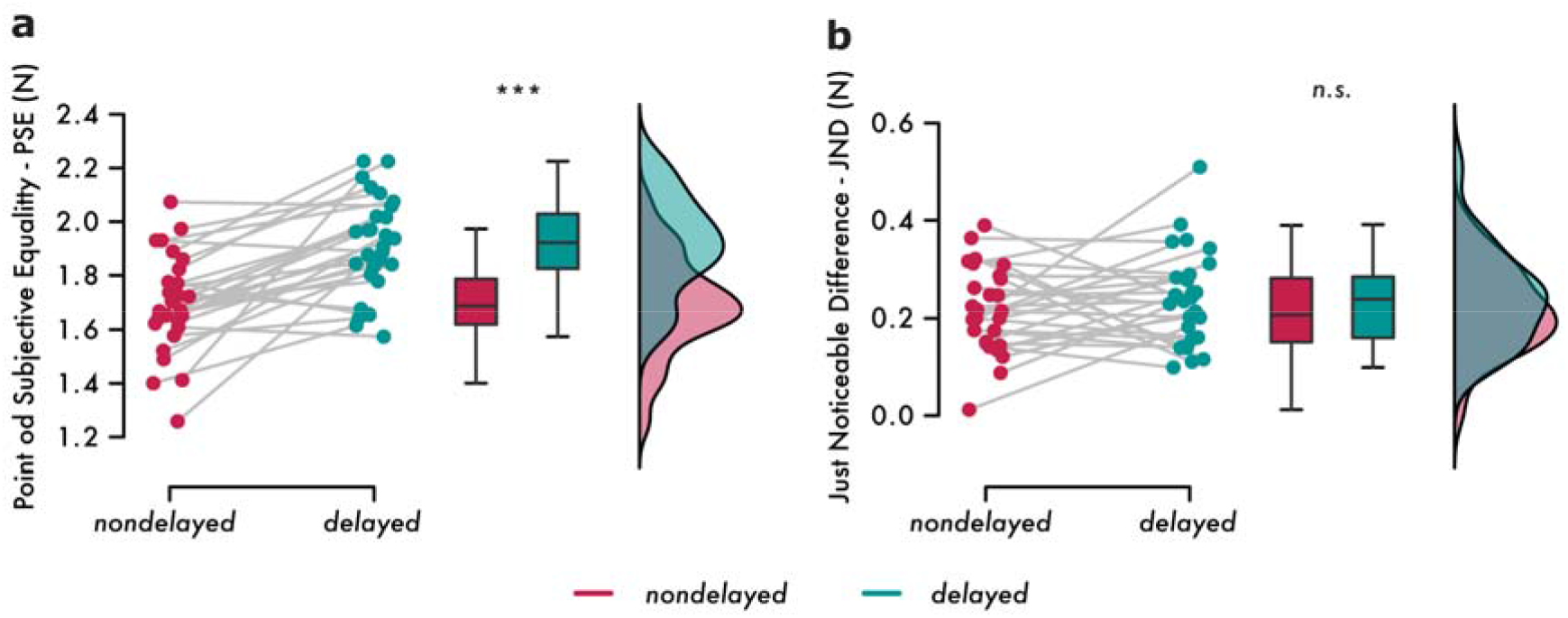
Results from the psychophysics task. **(a)** Individual PSEs and line plots illustrating the decreased PSE in the *nondelayed* compared to the *delayed self-generated touch* condition (*p* < 0.001). Boxplots and raincloud plots illustrate the group effects. **(b)** Individual JNDs and line plots illustrate the non-statistically significant JND changes between the *nondelayed* and the *delayed self-generated touch* conditions. Boxplots and raincloud plots illustrate the group effects.

### No changes in psychophysical responses evoked by temporal perturbations over time

As we expected, and in agreement with our previous results (Kilteni et al., 2019), we found no evidence for learning of the injected delay between the responses on the first and the second half of the psychophysical task (*n* = 28, *t*(27) *=* 0.418, *p* = 0.679, *Cohen’s d* = 0.079, *CI*^*95*^ = [−0.099, 0.149]). A Bayesian analysis also provided evidence for the absence of a learning effect (*BF*_*01*_ = 4.602) (**Supplementary Figure S2**).

### Temporal perturbations disrupt the attenuation of somatosensory reafference in the right primary and secondary somatosensory cortices and the right cerebellum

Compared to the baseline (*rest* blocks), both *nondelayed* and *delayed self-generated touch* conditions elicited significant neural activity (*p* < 0.05 *FWE-corrected*), including the contralateral premotor and motor cortices, supplementary motor area, and bilateral somatosensory and cerebellar areas, as expected (**Supplementary Tables S1** and **S2, Figure S3**). Importantly, when directly contrasting the two conditions, the *delayed self-generated touch* elicited increased activity in the right primary somatosensory cortex (right S1, postcentral gyrus) (MNI: *x* = 48, *y* = −18, *z* = 60; *p* = 0.002 *FWE-corrected*; *x* = 50, *y* = −16, *z* = 56; *p* = 0.002 *FWE-corrected*), and the right secondary somatosensory cortex (right SII, parietal operculum) (MNI: *x* = 42, *y* = −20, *z* = 16; *p* = 0.006 *FWE-corrected*) compared to the *nondelayed self-generated touch* condition (**Figure 4a-d, Supplementary Table S3**). Moreover, the *delayed self-generated touch* condition elicited increased activity in the right cerebellum (right CB) (MNI: *x* = 36, *y* = −72, *z* = −34; *p* = 0.049 *FWE-corrected*) compared to the *nondelayed self-generated touch* condition (**Figure 4e-f**) in lobule VIIa Crus I. No significant differences were observed in the hemispheres of other cerebellar lobules. The opposite contrast (*nondelayed* > *delayed* self-generated touch) mainly revealed activity in the right middle frontal gyrus that did not survive corrections for multiple comparisons and will therefore not be considered further (**Supplementary Figure S4, Supplementary Table S4**). Together, these findings show that a self-generated touch elicits stronger somatosensory and cerebellar activity when delivered with a 100 ms delay compared to an identical self-generated touch delivered at the time of contact between the two fingers. In other words, the neural attenuation of somatosensory reafference in the somatosensory and cerebellar cortices observed when the touch is delivered at its expected time (*nondelayed*) gets disrupted when the same touch is delivered with a delay (*i*.*e*., 100 ms).

**Figure 4.**
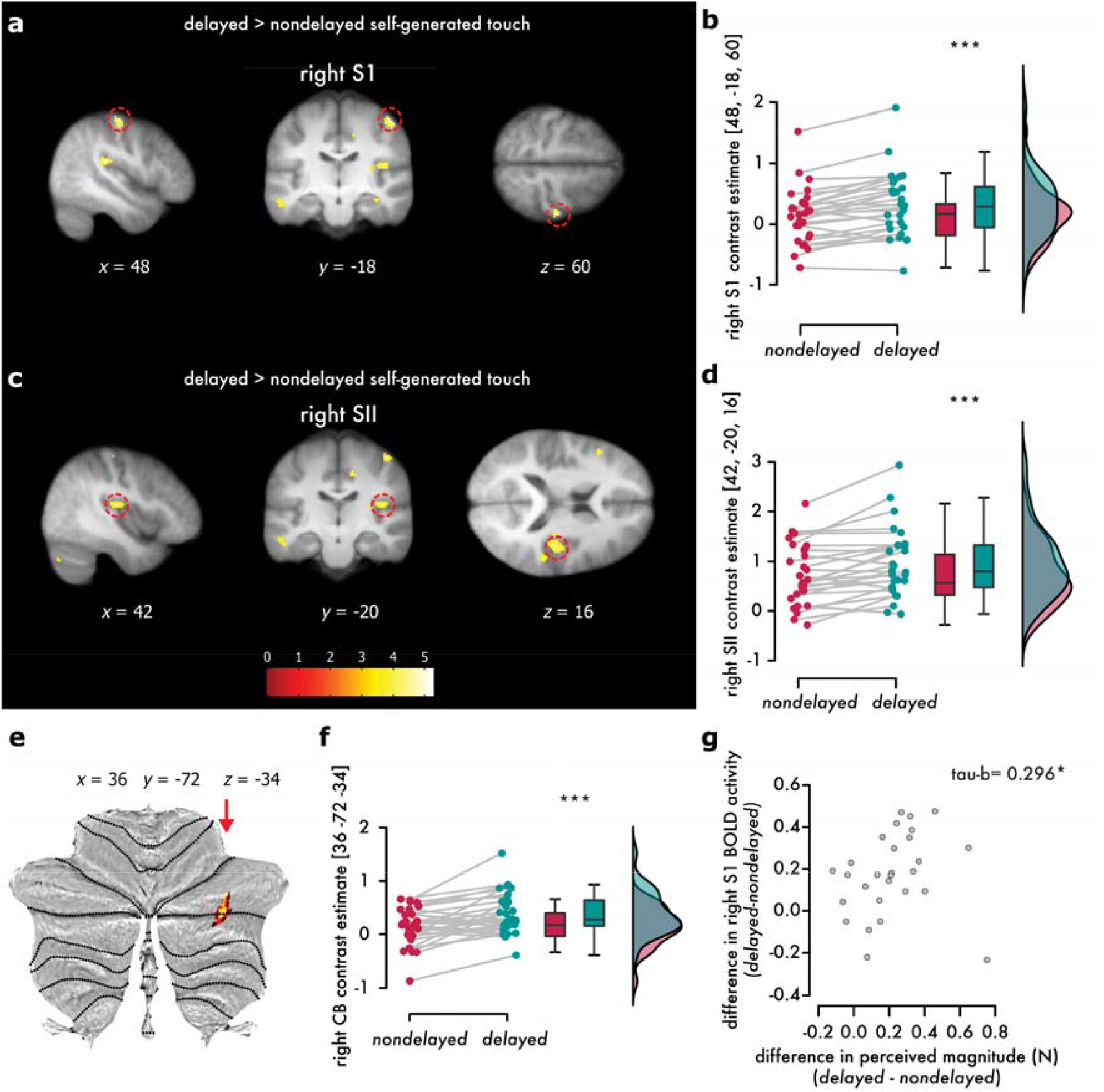
Somatosensory and cerebellar activations elicited during the *delayed* compared to the *nondelayed self-generated touch*. (**a, c**) Sagittal (left), coronal (middle), and axial (right) views of significant peaks of activation (*p < 0*.*05 FWE-corrected*) located at the right primary (postcentral gyrus) and secondary somatosensory cortex (parietal operculum). The activations maps have been rendered on the mean structural image across all 28 participants and are displayed at a threshold of *p* < 0.001 *uncorrected*. Red circles indicate the main significant peaks. (**e**) The cerebellar activations have been rendered on a flat representation of the human cerebellum (Diedrichsen and Zotow, 2015) at a threshold of *p* < 0.001 uncorrected. The red arrow indicates the significant peak within lobule VIIa Crus I (*p < 0*.*05 FWE-corrected*). (**b, d, f**) Individual contrast estimates and line plots illustrating the increase in the activation of the **(b)** right primary (rS1), **(d)** secondary somatosensory cortex (rSII), and (**f**) right cerebellum in the *delayed* compared to the *nondelayed self-generated touch* condition. All data have been corrected for multiple comparisons (*p* < 0.05 *FWE-corrected*). (**g**) Scatterplot showing the statistically significant and positive relationship between the difference in the perceived magnitude between the two conditions (*i*.*e*., difference in PSEs between the *delayed* and *nondelayed self-generated touch* conditions) and the difference in the BOLD activity of the right S1 between *delayed* and *nondelayed self-generated touch* conditions.

Including the forces generated by the right index finger (*active* taps) as a parametric modulator of each trial and testing its effect in BOLD activity, revealed significant activity in the left motor cortex (precentral gyrus) expanding to the left primary somatosensory cortex (postcentral gyrus), and bilateral cerebellum (lobules V, VI) (**Supplementary Figure S5, Supplementary Table S5**). That is, the intensity of the *active* taps the participants pressed with their right index finger parametrically modulated the activity in contralateral sensorimotor and bilateral cerebellar cortices. No modulation of the right somatosensory or motor cortex was detected (even at *p* < 0.005 *uncorrected*): *i*.*e*., the effects produced by the right hand’s force production and observed in the *left* hemisphere were anatomically distinct from the somatosensory effects in the *right* somatosensory cortex contralateral to the passive left index finger receiving the tactile stimulation (as reported above). Noteworthy, when including the parametric modulator in the main analysis contrasting the *delayed* and *nondelayed* self-generated touch conditions, we found the same somatosensory effects in the right S1 and right SII as the main analysis reported above (*delayed* > *nondelayed* self-generated touch (**Supplementary Table S6**). This rules out the possibility that small variations across trials in muscular contractions, produced force levels, or the associated somatosensory feedback from the right index finger, explain our main findings.

### The disruption of perceptual attenuation due to temporal perturbations predicts the disruption of the neural attenuation in the primary somatosensory responses

We then investigated whether the increase in PSEs due to the temporal perturbations (**Figure 3a**) was related to the increased responses of the right S1, right SII and right cerebellum (**Figure 4a, c, e**). To do so, we calculated the difference in the PSEs between the *delayed* and *nondelayed self-generated touch* conditions, and the difference in the contrast estimates for the activation peaks in the right S1 (MNI: *x* = 48, *y* = −18, *z* = 60), right SII (MNI: *x* = 42, *y* = −20, *z* = 16) and right cerebellum (MNI: *x* = 36, *y* = −72, *z* = −34) between the *delayed* and *nondelayed self-generated touch* conditions. The increase in PSEs significantly and positively predicted the increase in the responses of the right S1: *n* = 28, Kendall’s *tau-b* = 0.296, *p* = 0.027 (**Figure 4g**). No relationship was found for the right SII (*n* = 28, Pearson’s ρ = −0.083, *p* = 0.674) or the right cerebellum (*n* = 28, Pearson’s ρ = 0.1808, *p* = 0.2604). This suggests that the disruption of attenuation in the right S1 due to temporal perturbations reflects the disruption of attenuation at the perceptual level due to same temporal perturbations.

### No changes in the neural responses evoked by temporal perturbations over time

As we expected, and in agreement with our previous results and with the current psychophysical results reported above, we found no evidence for learning between the first and the last scanning block at the right S1 (*n* = 28, *t*(27) *=* 0.955, *p* = 0.348, *Cohen’s d* = 0.181, *CI*^*95*^ = [−0.323, 0.887]]), right SII (*n* = 28, *t*(27) *=* 0.670, *p* = 0.509, *Cohen’s d* = 0.127, *CI*^*95*^ = [−0.486, 0.956]), and right CB (*n* = 28, *t*(27) *=* 0.466, *p* = 0.645, *Cohen’s d* = 0.088, *CI*^*95*^ = [−0.469, 0.744]). A Bayesian analysis provided evidence for the absence of a learning effect (right S1: *BF*_*01*_ = 3.293, right SII: *BF*_*01*_ = 4.061, right CB: *BF*_*01*_ = 4.513) (**Supplementary Figure S6**).

### Temporal perturbations decrease the functional connectivity of the right primary somatosensory cortex with the left supplementary motor area, the bilateral cerebellum, and the left secondary somatosensory cortex, proportionally to the reduction in somatosensory perception

We expected that the disruption in the predictive processing of somatosensory reafference due to the perturbations should disrupt the connectivity of the right somatosensory cortices with brain areas involved in predicting the sensory consequences of the movement (*i*.*e*., SMA and cerebellum). To test this, we performed a seed-to-voxel generalized psychophysiological interaction (*gPPI*) analysis of functional connectivity using the right S1 or right SII as seeds and including the participants’ PSE differences from the psychophysical task (**Figure 3a**) as a covariate. This allowed us to isolate somatosensory functional connectivity increases or decreases that scaled linearly with the participants’ perceptual changes in their somatosensory perception. We found that the right S1 showed significant decreases in its connectivity with the left SMA (MNI: *x* = −2, *y* = −2, *z* = 52; *p* < 0.01 *FWE-corrected*) (**Figure 5a-b**), the left cerebellar lobule VIII (MNI: *x* = −30, *y* = −44, *z* = −58; *p* = 0.001 *FWE-corrected*; MNI: *x* = −16, *y* = −62, *z* = −60; *p* = 0.043 *FWE-corrected*) (**Figure 5c-d**), the right cerebellar lobule VIII (MNI: *x* = 22, *y* = −48, *z* = −58; *p* = 0.029 *FWE-corrected*; MNI: *x* = 20, *y* = −62, *z* = −60; *p* = 0.049 *FWE-corrected*) (**Figure 5e-f**), and the left secondary somatosensory cortex (MNI: *x* = 46, *y* = −16, *z* = 24; *p* = 0.023 *FWE-corrected*) during the *delayed* compared to the *nondelayed self-generated touch* condition (**Supplementary Table S7**). Similarly, the right SII showed a significant decrease in its connectivity with the left SMA (MNI: *x* = 22, *y* = −48, *z* = −58; *p* = 0.029 *FWE-corrected*) (**Figure 5g-h**). In contrast, there were no significant connectivity increases with the right S1 or SII as seeds. Together, these results indicate that the stronger was the disruption of the somatosensory reafference due to the temporal perturbation at the perceptual level (difference in PSEs), the weaker became the connectivity of the right S1 with the left SMA, the bilateral cerebellar lobules VIII, and the right SII, as well as the connectivity of the right SII with the left SMA.

**Figure 5.**
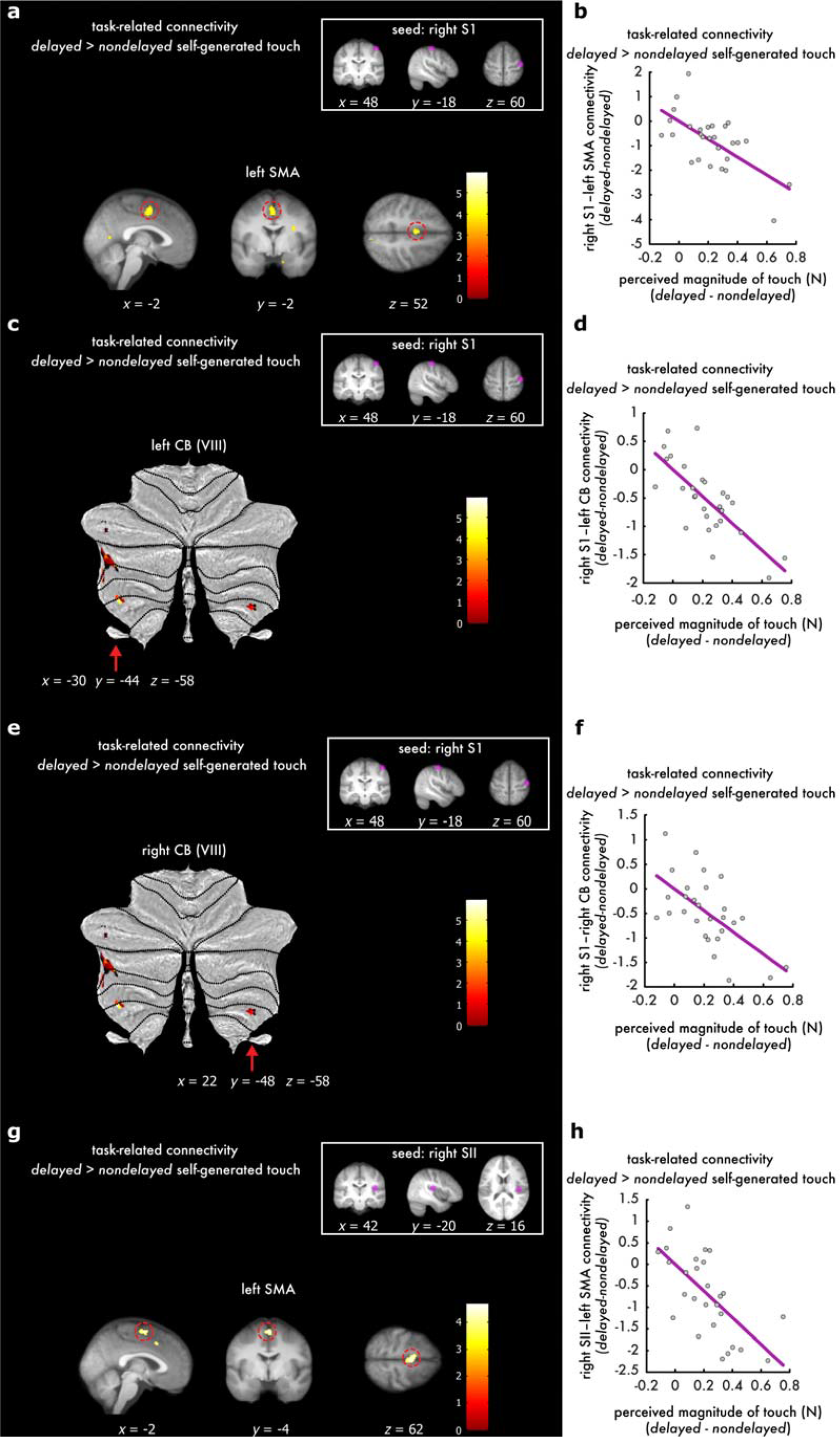
Functional connectivity results showing decreased connectivity of the left SMA and bilateral cerebellum with the right S1 (a-f) or right SII (g-h) as seeds, as a function of the participants’ somatosensory perception assessed psychophysically. **(a)** Sagittal (left), coronal (middle), and axial (right) views of the significant peak in the left SMA (*p < 0*.*05 FWE-corrected*) that decreased its connectivity with the right S1 (seed) in the generalized psychophysiological interaction analysis (gPPI). Activations have been rendered on the mean structural image across all participants at a threshold of *p* < 0.001 uncorrected and the red circles indicate the significant peak. **(c, e)** Cerebellar flatmaps showing the left **(c)** and right **(e)** cerebellar areas (VIII) that decreased their connectivity with the right S1. The cerebellar activations have been rendered on the cerebellar flatmap at a threshold of *p* < 0.001 uncorrected and red arrows indicate the location of the significant peaks *p < 0*.*05 FWE-corrected*). **(g)** Sagittal (left), coronal (middle), and axial (right) views of the significant peak in the left SMA (*p < 0*.*05 FWE-corrected*) that decreased its connectivity with the right SII (seed). Activations have been rendered on the mean structural image across all participants at a threshold of *p* < 0.001 uncorrected and the red circles indicate the significant peak (*p < 0*.*05 FWE-corrected*). **(b, d, f, h)** Scatterplots showing the relationship between the connectivity decreases between the corresponding seed and the significant peaks **(a, c, e, g)**, and the participants’ PSE differences extracted from the force-discrimination task.

### Temporal perturbations increase the functional connectivity of the left supplementary motor area with the left cerebellum

Finally, we hypothesized that the temporal perturbations should increase the connectivity of areas involved in motor planning (*i*.*e*., SMA) with areas processing the temporal discrepancy (*i*.*e*., cerebellum). Such connectivity changes could indicate the processing of the temporal error between the predicted and the actual somatosensory input to update the motor plan if needed. A seed-to-voxel functional connectivity analysis (*gPPI*) using the left SMA as the seed region confirmed this hypothesis: there were significant connectivity increases of the left SMA with the left cerebellum (lobules VI, VI/Crus I, VIIIa, VIIIb) (**Figure 6, Supplementary Table S8**) during the *delayed* compared to the *nondelayed self-generated touch* condition (*p* < 0.05 *FWE-corrected*). The effects did not covary with the participants’ perception, and no effects were found for the opposite contrast (*nondelayed* > *delayed self-generated touch*) (**Supplementary Table S9**).

**Figure 6.**
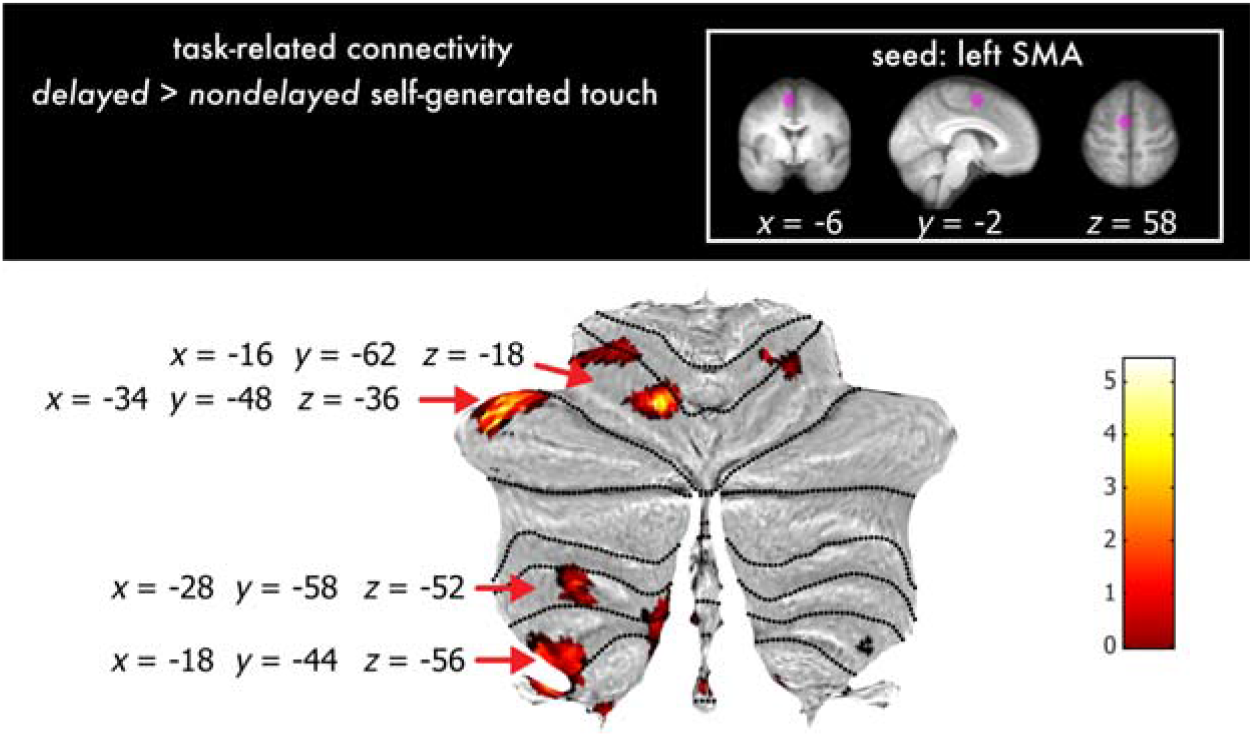
Functional connectivity results showing increased connectivity of the left SMA (seed) with the cerebellum during temporal perturbations. A generalized psychophysiological interactions analysis (*gPPI*) revealed multiple cerebellar peaks that increased connectivity with the left supplementary motor area during temporal perturbations compared to their absence (*delayed* > *nondelayed* self-generated touch). The cerebellar activations have been rendered on the cerebellar flatmap at a threshold of *p* < 0.001 uncorrected and red arrows indicate the location of the significant peaks within lobules VI, VI/VIIa, VIIIa and VIIIb.

## Discussion

Computational theories propose that internal forward models in the brain use information from our motor command to predict the timing of the sensory consequences of our movements and attenuate sensory input presented at that specific timing (Blakemore et al., 2000b; Wolpert and Flanagan, 2001; Bays and Wolpert, 2008). In contrast to previous neuroimaging studies imposing large temporal perturbations and thus contrasting somatosensory reafference with exafference conditions (Blakemore et al., 2001; Shergill et al., 2013), the present study focused on comparing conditions of identical somatosensory reafference with or without a brief temporal perturbation. This allowed us to test for the first time whether this time-locked predictive attenuation gets disrupted when temporal perturbations, as brief as of 100 ms, are introduced between the predicted and the actual timing of the somatosensory reafference.

At the perceptual level, we found that somatosensory reafference (*i*.*e*., self-generated touch) feels stronger when delivered with a 100 ms delay compared to when it is received at its predicted timing. These perceptual effects were mirrored at the neural level: both the right primary and secondary somatosensory cortices showed increased activity in presence of the temporal perturbations compared to their absence. Importantly, the disruption of perceptual attenuation was significantly correlated with the disruption of the neural attenuation of the right primary somatosensory cortex: that is, participants for whom the temporal perturbation had a larger effect in their perception, were the ones for whom the temporal perturbation had a larger effect in their somatosensory responses. Together, these results bring two novel conclusions. First, they demonstrate that somatosensory reafference is attenuated at both primary and secondary somatosensory cortex, in contrast to previous studies reporting effects only at the secondary somatosensory cortex when contrasting somatosensory reafference with exafference (Blakemore et al., 1998; Kilteni and Ehrsson, 2020). Given that SI is considered to be the earliest processing node in the cortical somatosensory processing system (Kandel et al., 2000), and that SII receives information from S1 through ipsilateral corticocortical connections, our findings reveal that the sensorimotor prediction has an effect on somatosensory processing earlier than previously thought. Second, they reveal for the first time a direct relationship between perceptual and neural attenuation and suggest that the primary somatosensory cortex reflects the degree to which participants perceived the somatosensory reafference, even though touches had identical intensity (2 N) in both conditions.

In our univariate analysis, we observed increased activity in the right cerebellum during temporal perturbations, in agreement with earlier PET findings (Blakemore et al., 2001), but not in the left hemisphere. At first, the absence of left cerebellar activation seems puzzling given that the cerebellum contains ipsilateral body representations, and temporal perturbations relate to the touch applied on the left hand. A possible explanation can be the small size of the delay we injected during temporal perturbations, but this is unlikely since Shergill et al. (2013) imposed a longer delay of 500 ms and they did not observe left cerebellar activity either. Interestingly, a recent metanalysis on the robustness of cerebellar activation during visual and auditory sensorimotor errors including temporal perturbations, failed to detect consistent cerebellar activations across the examined studies (Johnson et al., 2019). The authors noticed that cerebellar activations were most prominent in experiments where participants adapted to the imposed perturbation. In relation, in one of our previous studies (Kilteni et al., 2019) we showed that when repeatedly exposed to delays in somatosensory reafference, participants learn to predict the delayed touch and start to attenuate it. On the contrary, in the present study we purposefully included few exposure trials to avoid such learning, and indeed both behavioral and univariate control analyses showed that a short exposure to delays did not produce any significant learning of the injected perturbation. Therefore, we speculate that this lack of adaptation can potentially explain the absence of left cerebellar effects.

Our functional connectivity analysis showed that the right primary somatosensory cortex decreased its connectivity with the supplementary motor area, the cerebellum, and the secondary somatosensory cortex during the temporal perturbations. Critically, this connectivity decrease was a function of the perceived amplitude of the touch: that is, participants for whom the temporal perturbation had a larger effect in their perception, were the ones for whom the temporal perturbation produced a larger decrease in their somatosensory connectivity with the other areas. Previous results contrasting somatosensory reafference with exafference reported an increased connectivity between cerebellum and somatosensory cortices during (nondelayed) self-generated input compared to externally generated input as a function of the participants’ perception (Kilteni and Ehrsson, 2020): stronger attenuation of self-generated touches compared to externally generated ones yielded stronger somatosensory connectivity with the cerebellum during self-generated touches compared to externally generated ones. The present findings extend these previous results in pure conditions of somatosensory reafference, and show that a 100 ms temporal perturbation injected in somatosensory reafference is sufficient to disrupt the corticocerebellar connectivity previously suggested to implement somatosensory attenuation (Kilteni and Ehrsson, 2020). This points to the remarkable temporal precision for sensorimotor predictions; a brief temporal error of 100 ms between the predicted and the actual sensory reafference produces similar disruption in somatosensory attenuation as unpredicted sensory exafference.

Our perceptual, neural, and somatosensory connectivity effects, when put together, are in strong agreement with the framework of an internal forward model predictively attenuating self-generated input (Wolpert and Flanagan, 2001; McNamee and Wolpert, 2019). Accordingly, the left premotor cortices generate the right hand’s motor command and the associated efference copy that is used by the cerebellum to predict the sensory consequences of the action, including the touch on the left index finger. The cerebellar prediction is used to attenuate the received somatosensory activity. However, in presence of delays in receiving the sensory input, the somatosensory activity is not attenuated and thus, the received touch feels stronger. This is exactly what we observed in our psychophysics task and the univariate analysis. Moreover, the cerebellar prediction based on the efference copy about the timing of the sensory consequences precedes the delayed sensory feedback and this leads to weaker interaction with somatosensory areas. In line with this framework, our connectivity patterns showed a decrease in the connectivity between the primary and secondary somatosensory cortex (sensory feedback), cerebellum (forward model) and SMA (efference copy).

During the brief temporal perturbations, we observed that the SMA contralateral to the moving hand, increased its connectivity with the cerebellum (lobules VIII) during temporal perturbations. These findings assign for the first time a critical role to the SMA connectivity, for contrasting conditions of somatosensory reafference with and without subtle temporal perturbations. The SMA is target of cerebellar projections (Akkal et al., 2007; Bostan and Strick, 2018) and its posterior part (SMA proper) is connected to the corticospinal tract, precentral gyrus (M1), and ventrolateral thalamus (Johansen-Berg et al., 2004). Both SMA and cerebellum have been involved in temporal processing and temporal predictions (Rao et al., 1997; Ullén et al., 2003; Ivry and Schlerf, 2008; Wiener et al., 2010; Coull et al., 2011; Merchant and Yarrow, 2016) with the posterior SMA being particularly involved in sensorimotor sub-second temporal processing compared to the anterior SMA (Schwartze et al., 2012). The SMA is involved in motor planning and preparation (Makoshi et al., 2011; Ruan et al., 2018), and Transcranial Magnetic stimulation (TMS) over SMA during voluntary movements produces perceptual effects consistent with a disruption of the efference copy that allows the prediction and attenuation of somatosensory responses (Haggard and Whitford, 2004). Similarly, the cerebellum is considered to implement the forward model (Shadmehr et al., 2008; McNamee and Wolpert, 2019; Popa and Ebner, 2019), and cerebellar TMS produces perceptual effects consistent with a disruption of the sensorimotor prediction and its combination with the actual sensory feedback (Miall et al., 2007). From a theoretical perspective, functional connectivity between SMA and cerebellum could refer to (a) the efference copy being sent to the cerebellar forward model to predict the sensory consequences of the movement, but also (b) to the error signal sent back to the SMA to inform the motor centers for the errors. Our connectivity analysis cannot distinguish between these two scenarios. However, given that the sensorimotor efference copy based predictions should be computed independently of temporal perturbations, and that this connectivity *increased* during temporal perturbations, we propose that the most compatible interpretation is that of communicating the temporal prediction error.

Disturbances in attenuating somatosensory reafference have been repeatedly reported in patients with schizophrenia (Blakemore et al., 2000a; Shergill et al., 2005, 2014) and non-clinical individuals with high schizotypal personality traits (Asimakidou et al., 2022). Using encephalography, it was further shown that schizophrenic patients suppress their nondelayed self-generated sounds to a lesser extent compared to healthy controls, but show normal attenuation when the auditory reafference is delayed by a 50 or 100 ms delay (Whitford et al., 2011). We therefore speculate that the pattern of effects revealed by the present study might reverse for such patients, leading to the attenuation of the delayed somatosensory reafference but not the nondelayed one – a speculation that should be tested in future experiments.

## Author Contributions

K.K. and H.H.E. conceived and designed the experiment. K.K. and C.H. collected the data. K.K conducted the statistical analysis. K.K. and H.H.E. wrote the manuscript and C.H. approved the final version of the manuscript.

## Acknowledgments

Konstantina Kilteni was supported by the Swedish Research Council (VR Starting Grant 2019-01909 granted to K.K.) and the Marie Skłodowska-Curie Intra-European Individual Fellowship (#704438). The project was funded by the Swedish Research Council, Torsten Söderbergs Stiftelse, and Göran Gustafssons Stiftelse.

## Supplementary material

**Figure S1.**
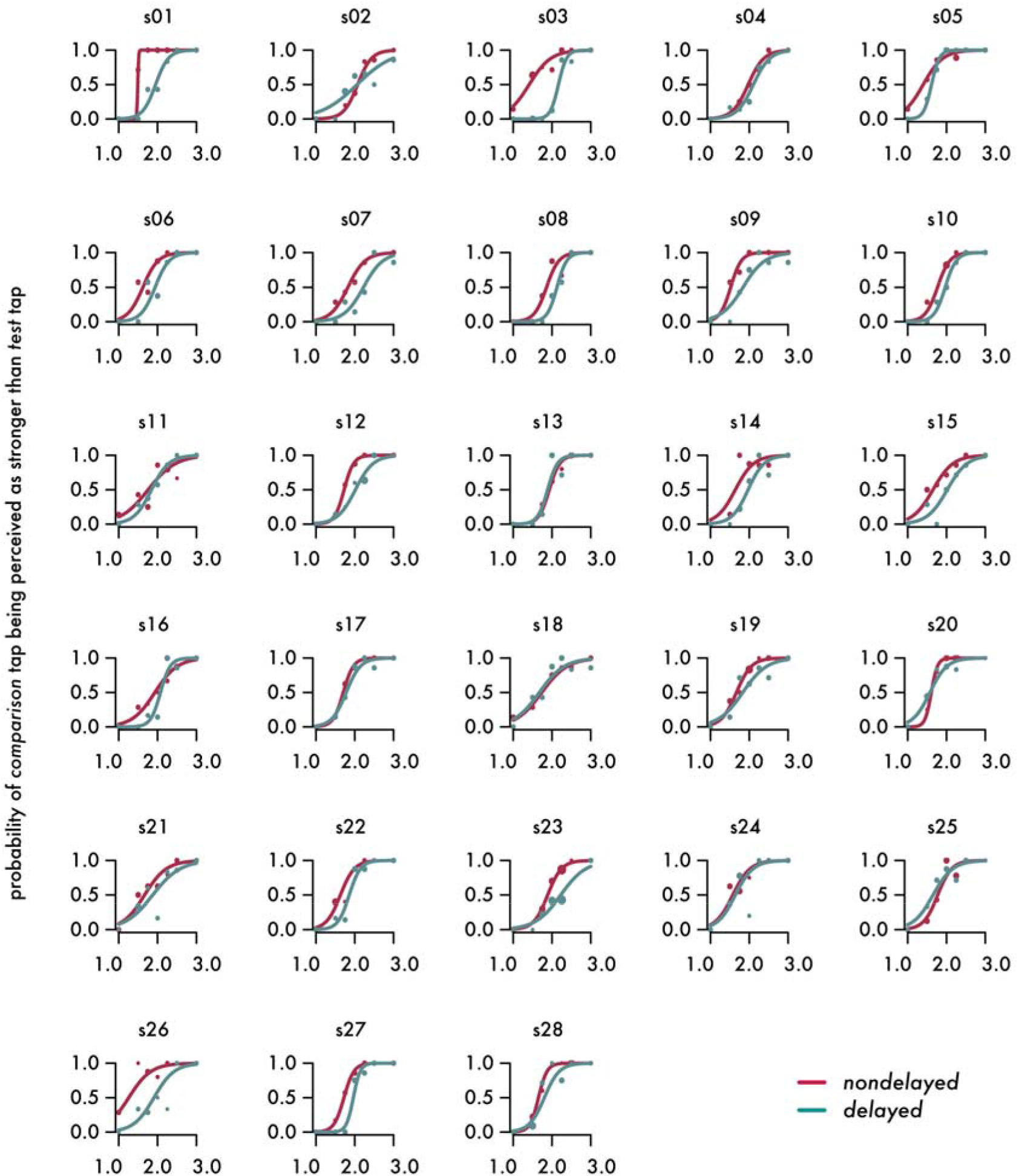
Individual plots for the psychophysical session. The marker size is proportional to the number of repetitions for stimulus level. For all participants and conditions, the fitted model resulted to a McFadden’s R squared measure ranging between 0.409 and 0.945.

**Figure S2.**
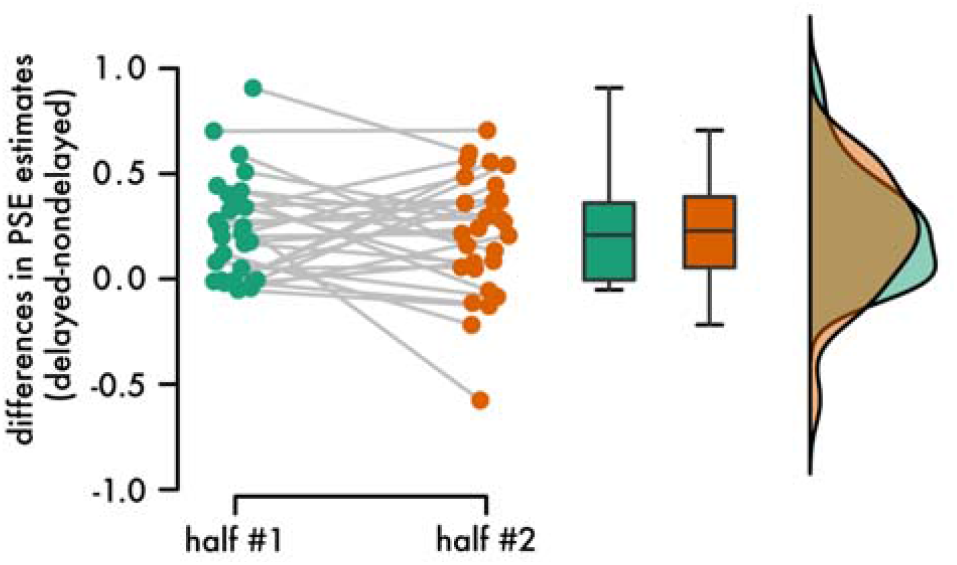
Absence of learning effects of the 100 ms delay during the psychophysical session. Individual differences and line plots illustrating the difference in the PSEs between the two conditions (*delayed* – *nondelayed* self-generated touch) for the first and the second half of the psychophysical task. Boxplots and raincloud plots illustrate the group effects. There were no learning effects, as strongly supported by a Bayesian analysis.

**Table S1.**
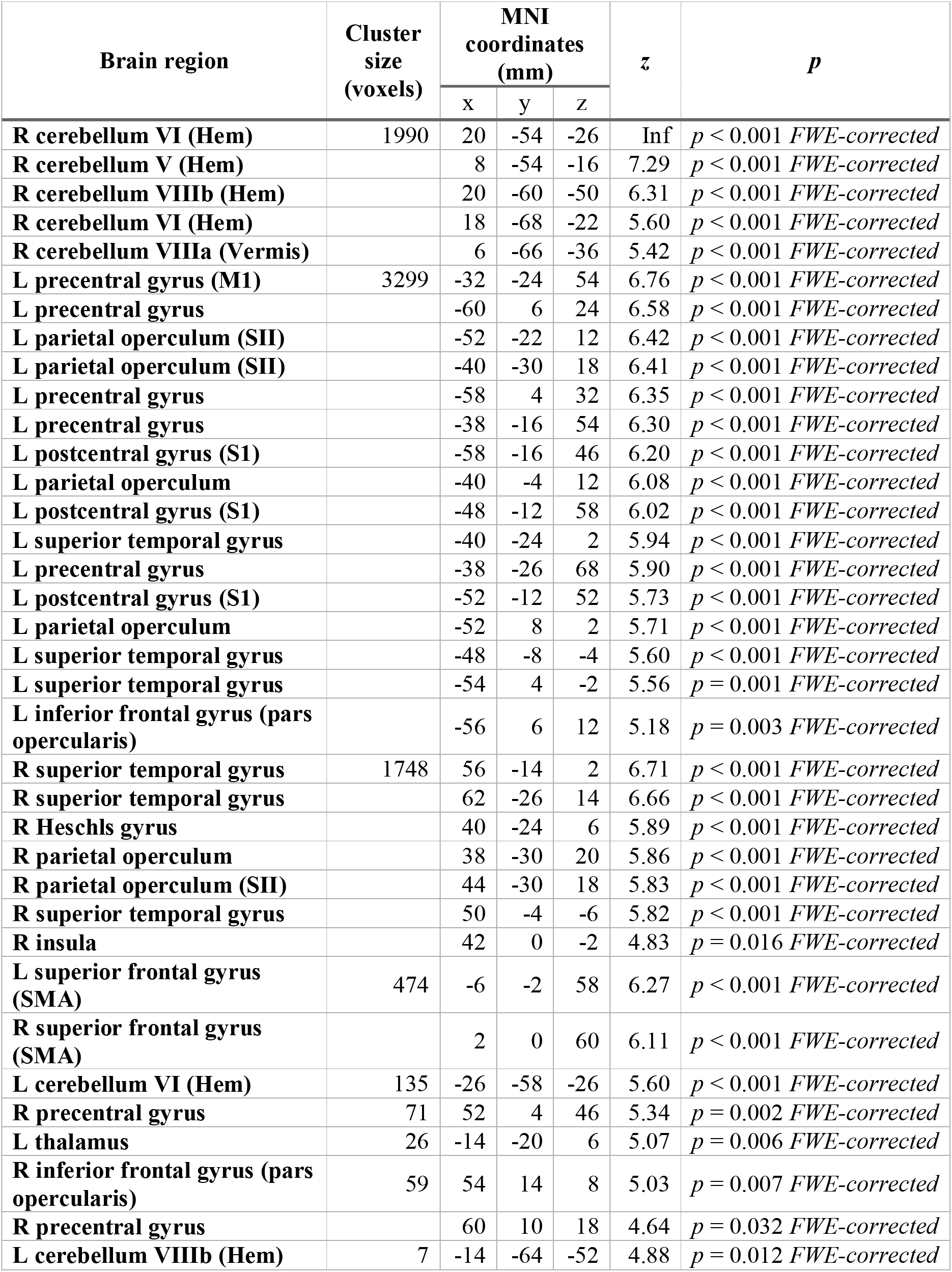
Activation peaks for the *nondelayed self-generated touch*. Peaks reflecting greater effects during *nondelayed self-generated touch* compared to rest (*nondelayed* > 0). Only the peaks that survived the FWE correction (*p* < 0.05) belonging to clusters with size greater than 4 voxels are reported for spatial restrictions.

**Table S2.**
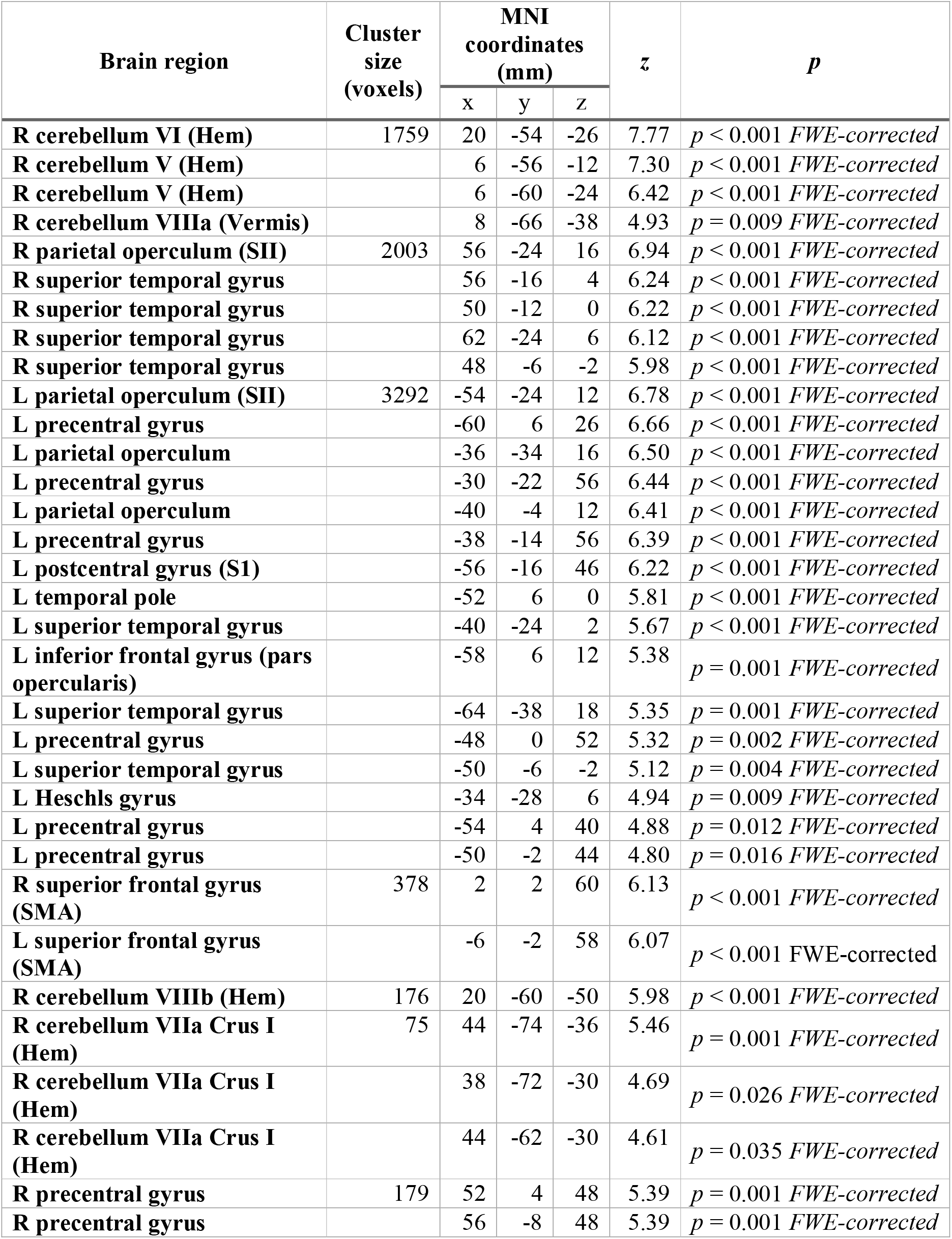
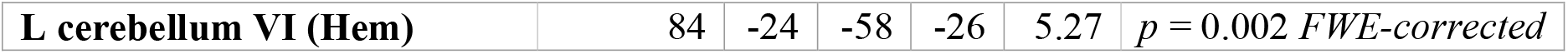
Activation peaks for the *delayed self-generated touch*. Peaks reflecting greater effects during *delayed self-generated touch* compared to rest (*delayed* > 0). Only the peaks that survived the FWE correction (*p* < 0.05) belonging to clusters with size greater than 4 voxels are reported for spatial restrictions.

**Figure S3.**
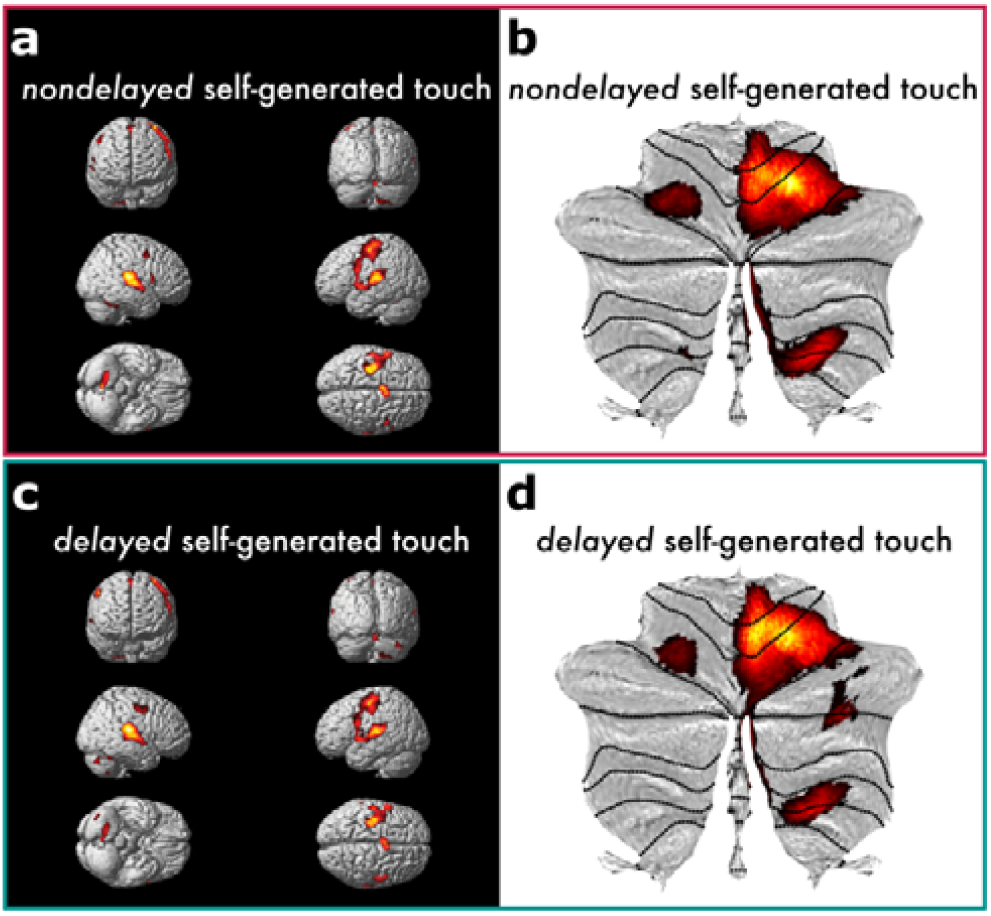
Activations during *nondelayed* and *delayed* self-generated touch. **(a-b)** Activations reflect greater effects during *nondelayed* self-generated touch compared to rest. **(c-d)** Activations reflect greater effects during *delayed* self-generated touch compared to rest. In both contrasts, auditory areas were also activated since the participants heard auditory GO cues to produce the self-generated touches. **(a, c)** The activations have been rendered on the standard single subject 3D-volume provided with SPM. **(b, d)** Cerebellar activations have been overlaid onto a cerebellar flatmap. **(a-d)** All activation maps are displayed at a threshold of p < 0.05 FWE-corrected.

**Table S3.** Activations that are greater during the *delayed* than the *nondelayed self-generated touch* conditions. Peaks reflecting greater effects during *delayed* compared to *nondelayed self-generated touch* (*delayed* > *nondelayed*).

| Brain region | Cluster size (voxels) | MNI coordinates (mm) |  |  | z | p |
| --- | --- | --- | --- | --- | --- | --- |
|  |  | x | y | z |  |  |
| **R parietal operculum** | 86 | 50 | -30 | 20 | 4.33 | $p < 0.001$ *uncorrected* |
| **R postcentral gyrus (S1)** | 59<sup>1</sup> | 48 | -18 | 60 | 4.07 | $p = 0.002$ *FWE-corrected* |
| **R postcentral gyrus (S1)** | | 50 | -16 | 56 | 3.98 | $p = 0.002$ *FWE-corrected* |
| **R precentral/postcentral gyrus** | | 54 | -12 | 46 | 3.69 | $p < 0.001$ *uncorrected* |
| **L superior frontal gyrus** | 72 | -12 | 44 | 30 | 3.99 | $p < 0.001$ *uncorrected* |
| **L inferior frontal gyrus (pars orbitalis)** | 39 | -46 | 34 | -10 | 3.72 | $p < 0.001$ *uncorrected* |
| **L inferior frontal gyrus (pars triangularis)** | 91 | -40 | 22 | 22 | 3.72 | $p < 0.001$ *uncorrected* |
| **L inferior frontal gyrus (pars triangularis)** | | -48 | 20 | 16 | 3.38 | $p < 0.001$ *uncorrected* |
| **L inferior frontal gyrus (pars triangularis)** | | -54 | 26 | 10 | 3.24 | $p = 0.001$ *uncorrected* |
| **R parietal operculum (SII)** | 54<sup>2</sup> | 42 | -20 | 16 | 3.71 | $p = 0.006$ *FWE-corrected* |
| **R hippocampus** | 12 | 36 | -16 | -16 | 3.64 | $p < 0.001$ *uncorrected* |
| **R cerebellum VIIa Crus I (Hem)** | 49 | 36 | -72 | -34 | 3.57 | $p = 0.049$ *FWE-corrected* |
| **R middle cingulate gyrus** | 11 | 14 | -20 | 46 | 3.52 | $p < 0.001$ *uncorrected* |
| **L middle frontal gyrus** | 24 | -34 | 12 | 46 | 3.48 | $p < 0.001$ *uncorrected* |
| **L middle temporal gyrus** | 30 | -54 | -18 | -18 | 3.44 | $p < 0.001$ *uncorrected* |
| **L inferior parietal lobule** | 11 | -32 | -72 | 42 | 3.36 | $p < 0.001$ *uncorrected* |
| **L supramarginal gyrus** | 14 | -48 | -46 | 26 | 3.35 | $p < 0.001$ *uncorrected* |
| **R postcentral gyrus** | 8 | 60 | -2 | 36 | 3.32 | $p < 0.001$ *uncorrected* |
| **R insula** | 12 | 30 | -20 | 16 | 3.31 | $p < 0.001$ *uncorrected* |

\* After small volume correction.
<sup>1</sup> The cluster size is 106 before corrections for multiple comparisons and is reduced to 59 after small volume correction.
<sup>2</sup> The cluster size is 74 before corrections for multiple comparisons and is reduced to 54 after small volume correction.

**Figure S4.**
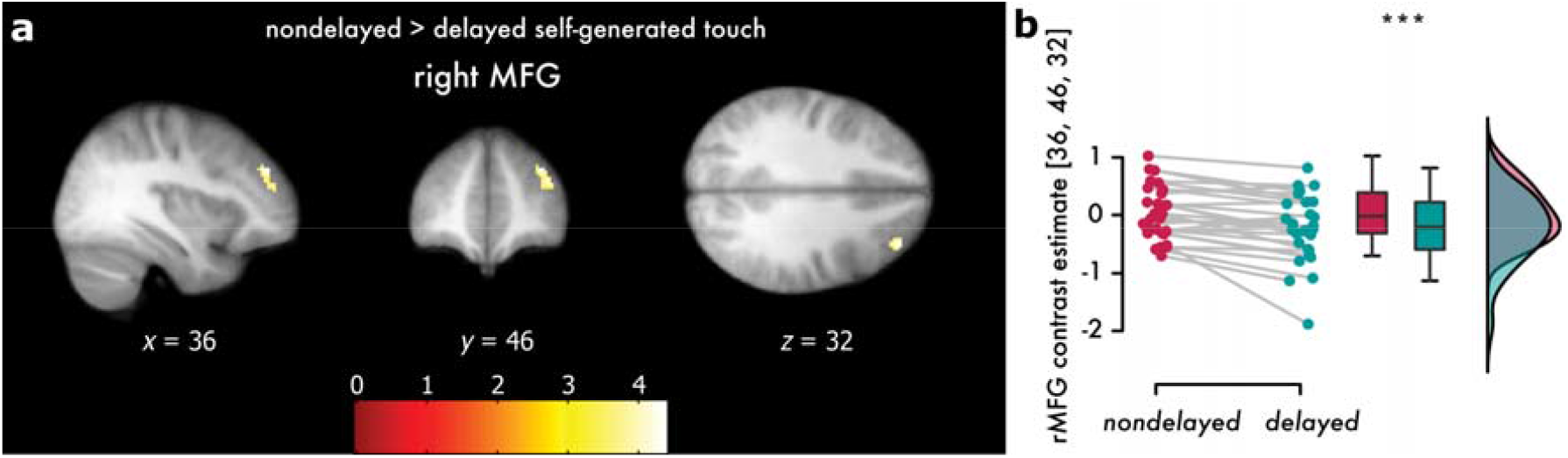
Activations elicited during the *nondelayed* compared to the *delayed self-generated touch*. (**a**) Activations reflect greater effects during *nondelayed* compared to *delayed self-generated touch* at the right middle frontal gyrus that did not survive corrections for multiple comparisons. The activations have been rendered on the mean structural image across all participants. All activation maps are displayed at a threshold of *p* < 0.001 uncorrected (**Supplementary Table S4**). (**b**) Individual contrast estimates and line plots illustrating the increase in the activation of the middle frontal gyrus in the *nondelayed* compared to the *delayed self-generated touch* condition (*p* < 0.001).

**Table S4.**
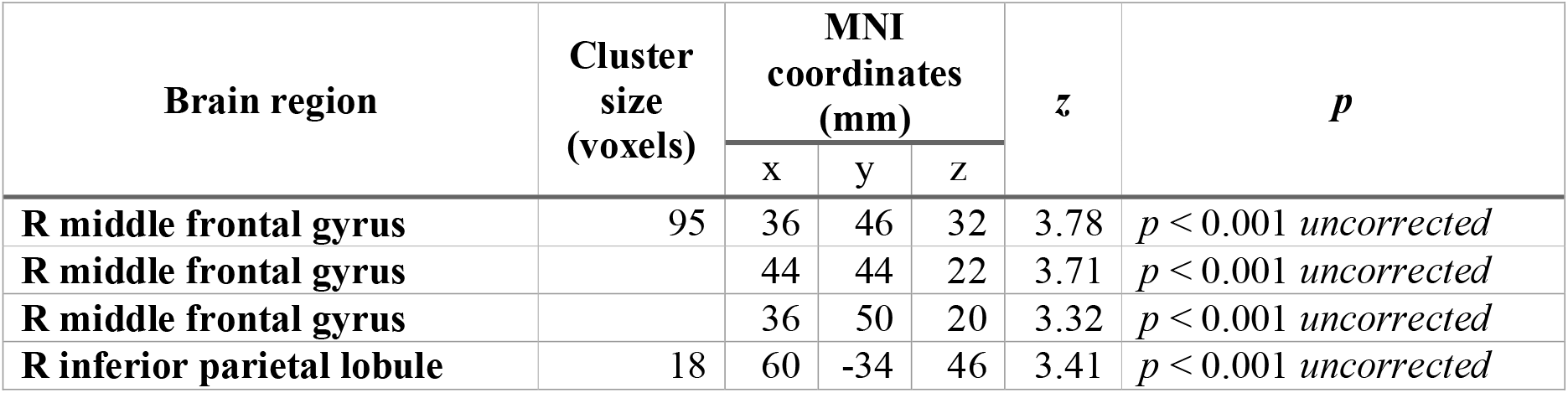
Activations that are greater during the *nondelayed* than the *delayed self-generated touch* conditions. Peaks reflecting greater effects during *nondelayed* compared to *delayed self-generated touch* (*nondelayed* > *delayed*).

| Brain region | Cluster size (voxels) | MNI coordinates (mm) |  |  | *z* | *p* |
| --- | --- | --- | --- | --- | --- | --- |
|  |  | x | y | z |  |  |
| **R middle frontal gyrus** | 95 | 36 | 46 | 32 | 3.78 | *p* < 0.001 *uncorrected* |
| **R middle frontal gyrus** |  | 44 | 44 | 22 | 3.71 | *p* < 0.001 *uncorrected* |
| **R middle frontal gyrus** |  | 36 | 50 | 20 | 3.32 | *p* < 0.001 *uncorrected* |
| **R inferior parietal lobule** | 18 | 60 | -34 | 46 | 3.41 | *p* < 0.001 *uncorrected* |

**Table S5.**
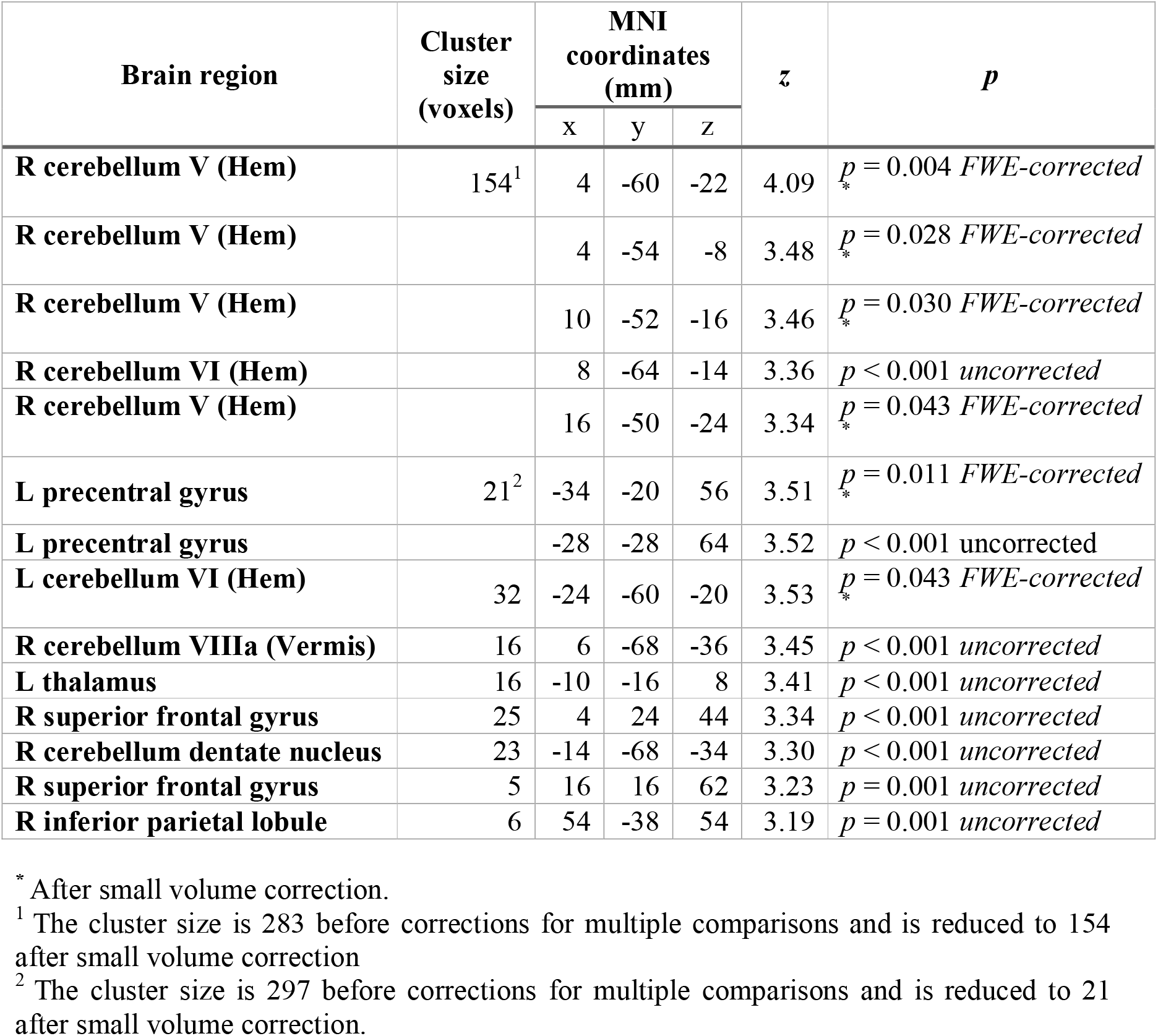
Areas whose activity was parametrically modulated by the strength of the right hand’s *active* taps across both *nondelayed* and *delayed* self-generated touch conditions.

**Figure S5.**
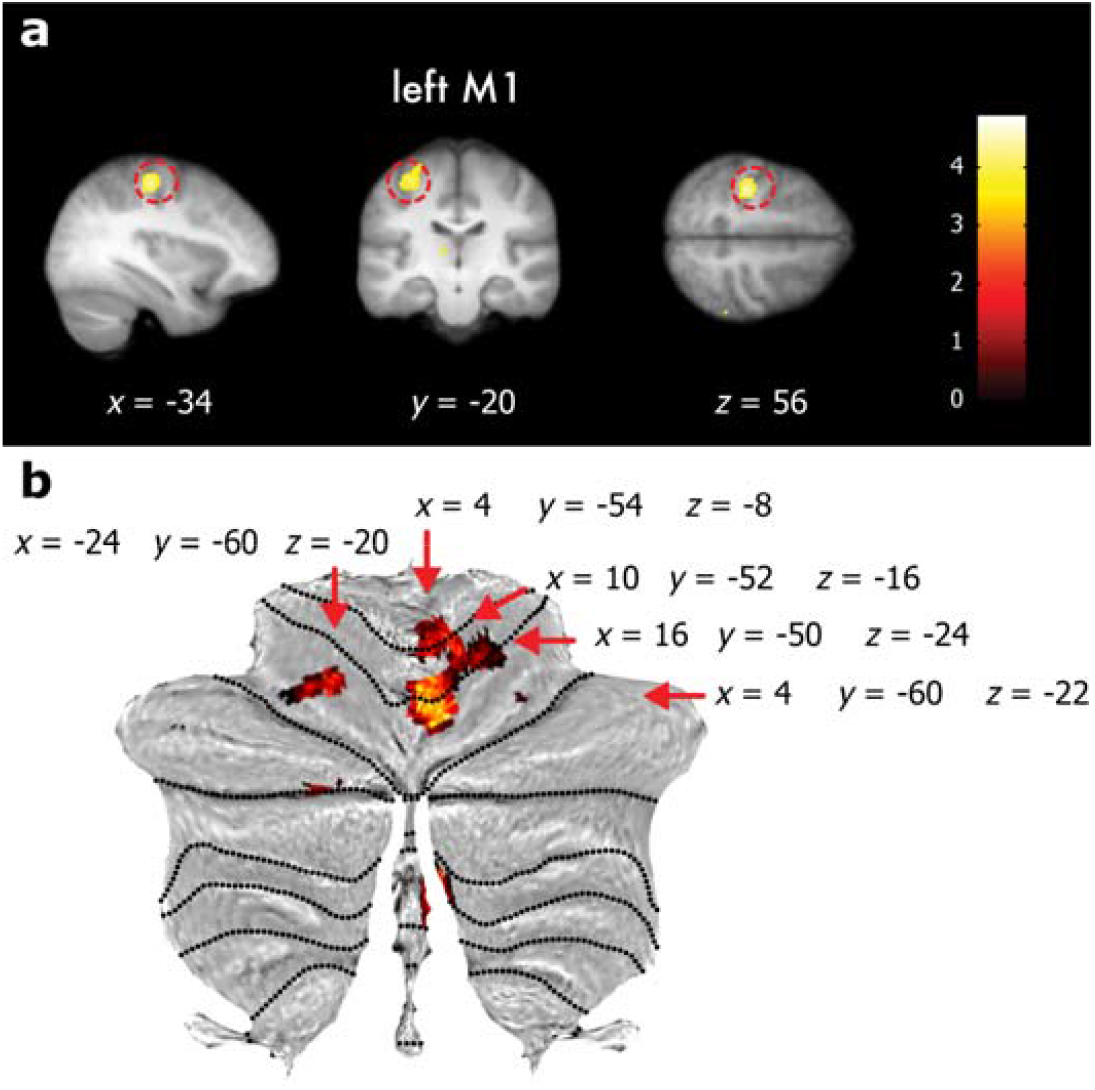
Sensorimotor (a) and cerebellar (b) areas whose BOLD activity was significantly and linearly modulated by the forces participants exerted with their right index finger (*active* taps). **(a)** The activity of the left motor cortex was significantly modulated by the strength of the *active* taps. The cluster extends to the left primary somatosensory cortex. The red circle indicates the significant peak. The activations have been rendered on the mean structural image across all participants at a threshold of *p* < 0.001 *uncorrected*. **(b)** Multiple peaks at the right and left cerebellum were significantly modulated by the taps of the participants’ right hand (lobule V, VI). Pointers denote the significant peaks. The cerebellar activations have been rendered on the cerebellar flatmap at a threshold of *p* < 0.001 *uncorrected*.

**Table S6.** Activations that are greater during the *delayed* than the *nondelayed self-generated touch* conditions, with *active* taps as parametric modulator. Peaks reflecting greater effects during *delayed* compared to *nondelayed self-generated touch* (*delayed* > *nondelayed*).

| Brain region | Cluster size (voxels) | MNI coordinates (mm) |  |  | z | p |
| --- | --- | --- | --- | --- | --- | --- |
|  |  | x | y | z |  |  |
| **R parietal operculum** | 222 | 50 | -30 | 20 | 4.62 | $p = 0.037$ FWE-corrected |
| **R parietal operculum (SII)** | | 42 | -18 | 16 | 3.73 | $p_* = 0.006$ FWE-corrected |
| **R postcentral gyrus (S1)** | 73<sup>1</sup> | 50 | -16 | 54 | 4.07 | $p_* = 0.002$ FWE-corrected |
| **R postcentral gyrus (S1)** | | 48 | -18 | 60 | 3.87 | $p_* = 0.002$ FWE-corrected |
| **L superior frontal gyrus** | 69 | -12 | 44 | 30 | 4.05 | $p < 0.001$ uncorrected |
| **R hippocampus** | 14 | 36 | -16 | -16 | 3.69 | $p < 0.001$ uncorrected |
| **L inferior frontal gyrus (pars triangularis)** | 73 | -40 | 22 | 22 | 3.67 | $p < 0.001$ uncorrected |
| **L inferior frontal gyrus (pars triangularis)** | | -48 | 20 | 16 | 3.34 | $p < 0.001$ uncorrected |
| **L inferior frontal gyrus (pars orbitalis)** | 30 | -48 | 34 | -8 | 3.62 | $p < 0.001$ uncorrected |
| **R cerebellum VIIa Crus I (Hem)** | 45 | 36 | -72 | -34 | 3.53 | $p_* = 0.054$ FWE-corrected |
| **R middle cingulate cortex** | 14 | 14 | -20 | 46 | 3.49 | $p < 0.001$ uncorrected |
| **L inferior parietal lobule** | 22 | -32 | -72 | 42 | 3.48 | $p < 0.001$ uncorrected |
| **L middle frontal gyrus** | 17 | -34 | 12 | 46 | 3.40 | $p < 0.001$ uncorrected |
| **R postcentral gyrus** | 9 | 60 | -2 | 36 | 3.35 | $p < 0.001$ uncorrected |
| **R insula** | 17 | 30 | -18 | 14 | 3.35 | $p < 0.001$ uncorrected |
| **L middle temporal gyrus** | 15 | -56 | -18 | -18 | 3.32 | $p < 0.001$ uncorrected |
| **L inferior parietal lobule** | 6 | -48 | -44 | 24 | 3.24 | $p = 0.001$ uncorrected |

\* After small volume correction.
<sup>1</sup> The cluster size is 145 before corrections for multiple comparisons and is reduced to 73 after small volume correction.

**Figure S6.**
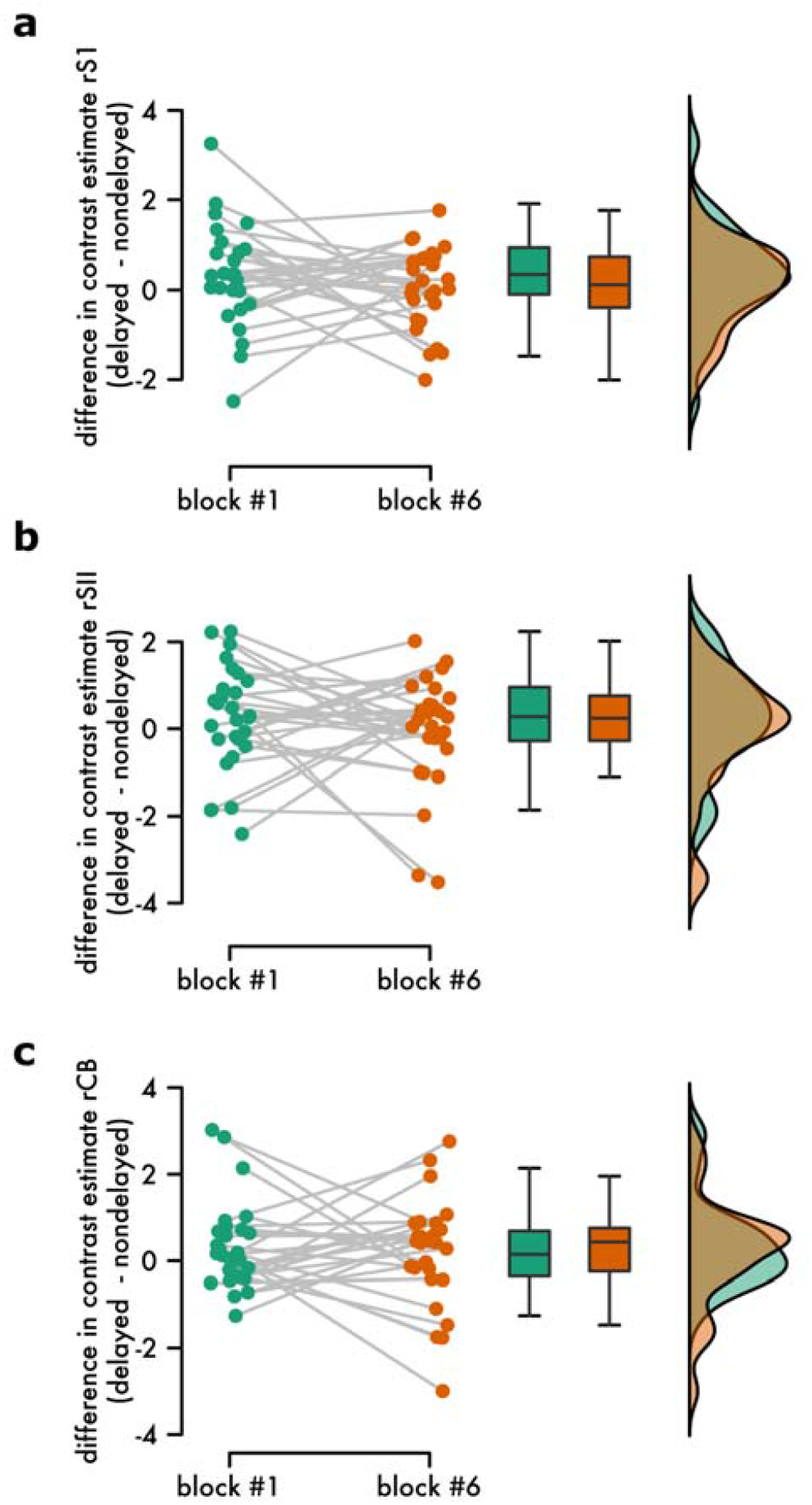
Absence of learning effects of the 100 ms delay during the fMRI run. Individual differences and line plots illustrating the difference in the extracted activity for contrast estimates between the two conditions (*delayed* – *nondelayed* self-generated touch) for the first and the last block of the fMRI run at the **(a)** right primary somatosensory cortex (rS1), **(b)** secondary somatosensory cortex (rSII), and **(c)** the right cerebellum (rCB). Boxplots and raincloud plots illustrate the group effects. There were no learning effects, as strongly supported by a Bayesian analysis.

**Table S7.**
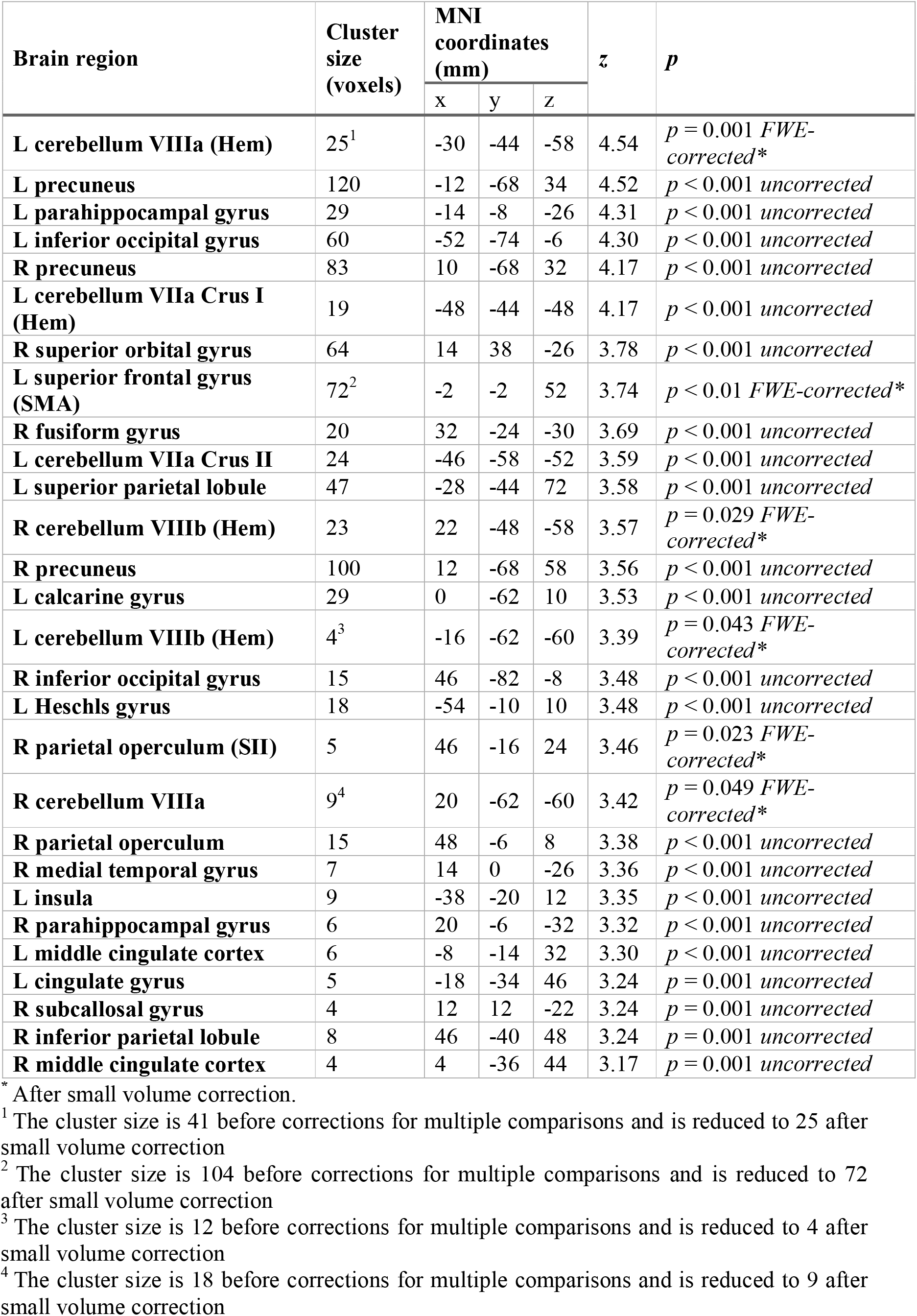
Peaks that decreased their connectivity with the left primary somatosensory cortex during the temporal mismatches as a function to the participants’ PSE difference between the *delayed* and *nondelayed self-generated touch* conditions. Peaks reflecting lower connectivity with the left primary somatosensory cortex when the touch was delayed compared to when it was nondelayed (*delayed* > *nondelayed*) and covaried with the participants’ perception. Only the peaks that belonged to clusters with size greater than 4 voxels are reported for spatial restrictions.

**Table S8.**
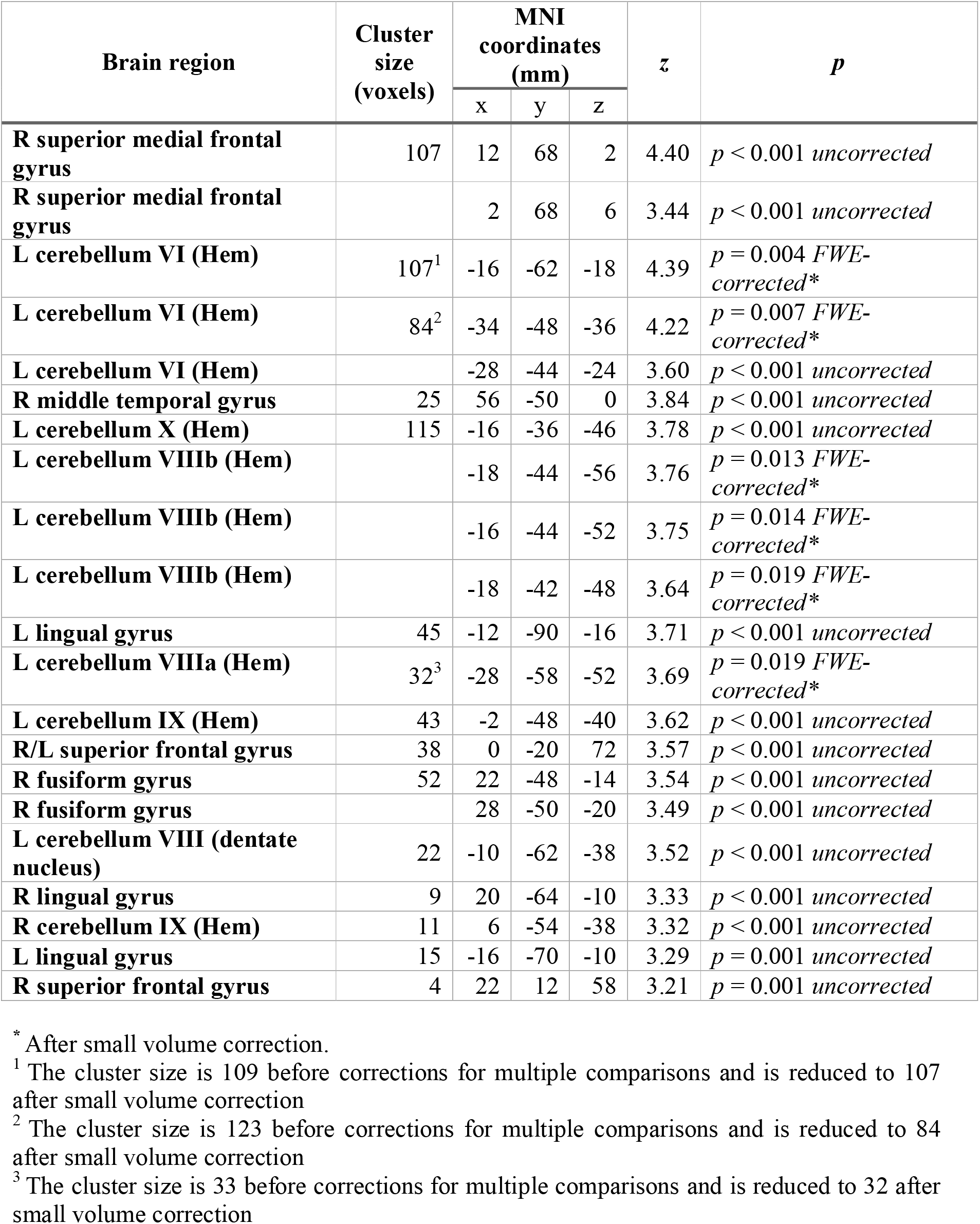
Peaks that increased their connectivity with the left supplementary motor area during the temporal mismatches. Peaks reflecting greater connectivity with the left supplementary motor area when the touch was delayed compared to when it was nondelayed (*delayed* > *nondelayed*). Only the peaks that belonged to clusters with size greater than 4 voxels are reported for spatial restrictions.

**Table S9.** Peaks that decreased their connectivity with the left supplementary motor area during the temporal mismatches. Peaks reflecting lower connectivity with the left supplementary motor area when the touch was delayed compared to when it was nondelayed (*delayed* > *nondelayed*). Only the peaks that belonged to clusters with size greater than 4 voxels are reported for spatial restrictions.

| Brain region | Cluster size (voxels) | MNI coordinates (mm) |  |  | z | p |
| --- | --- | --- | --- | --- | --- | --- |
|  |  | x | y | z |  |  |
| **R caudate nucleus** | 16 | 18 | 22 | 10 | 3.28 | $p = 0.001$ *uncorrected* |

